# A multi-omic atlas of human autonomic and sensory ganglia implicates cell types in peripheral neuropathies

**DOI:** 10.1101/2025.09.18.677119

**Authors:** Lite Yang, Adam J. Dourson, Ruichen Tao, Kevin Boyer, Pauline Meriau, John Del Rosario, Jiwon Yi, Richard A. Slivicki, Zachariah Bertels, Maria Payne, Juliet M. Mwirigi, Prashant Gupta, John Lemen, Bryan A. Copits, Guoyan Zhao, Valeria Cavalli, Alexander Chamessian, Robert W. Gereau

## Abstract

The human peripheral nervous system (PNS) consists of many ganglia including sympathetic ganglia (SG) and dorsal root ganglia (DRG). These ganglia house the cell bodies of diverse PNS neurons that transmit autonomic and sensory signals, as well as a much larger number of non-neuronal cells. However, the molecular and cellular diversity of these human PNS cell types and their implications in human disease remain elusive. By generating an integrated single-cell multi-omic atlas of human SG and DRG, we provide comprehensive transcriptional and epigenomic landscapes of various cell types in these peripheral ganglia. While the major cell types and their cell-type-specific transcriptional and epigenomic features are similar between human SG and DRG, we identify key differences between SG and DRG cell types. These differences highlight the distinct molecular and cellular mechanisms underlying their specific functions. We also profiled key genomic regulatory networks that govern cell-type-specific gene expression in these peripheral ganglia. Moreover, by mapping the expression and chromatin accessibility of disease-associated genes in human SG and DRG, we identify cell types that may underlie various peripheral neuropathies. This atlas serves as a valuable resource for understanding the intricate cell-type-specific molecules and interactions in the human PNS and their implications in human health and diseases.

## INTRODUCTION

The peripheral nervous system (PNS) includes the autonomic and somatic divisions. The autonomic division consists of the sympathetic arm, including the sympathetic ganglia (SG), and the parasympathetic arm. The somatic division comprises the sensory arm, including the dorsal root ganglia (DRG) and trigeminal ganglia, and the motor arm. These ganglia are distributed throughout the periphery and house the cell bodies of many neuron types and non-neuronal cells of the PNS. The SG is primarily involved in autonomic regulation, contributing to the “fight or flight” response and maintaining homeostasis, whereas the DRG plays a crucial role in sensory perception by relaying signals from the periphery to the central nervous system (CNS)[1, 2]. While SG and DRG differ in several key aspects, including innervation targets, neurotransmitter profiles, and physiological functions, they share a common developmental origin from the neural crest and encompass very similar cell types[3, 4].

Neurons in the SG, primarily noradrenergic with a smaller proportion of cholinergic neurons, innervate peripheral organs and tissues to regulate autonomic functions, such as cardiac output, blood glucose levels, body temperature, and immune responses[5]. In contrast, sensory neurons in the DRG are mostly glutamatergic and transmit information about mechanosensation, temperature, pain, and proprioception from the periphery to the CNS[1, 6-8]. Morphologically, sympathetic neurons are multi-polar, possessing an axon and multiple dendrites forming synapses with the preganglionic neurons from the spinal cord, whereas sensory neurons have bifurcating pseudo-unipolar axons, projecting one branch to the target organs (such as skin, viscera, and muscle) and the other to the CNS.

Glial cells, including satellite glial cells (or SGC) and Schwann cells, provide crucial support and regulation of the neuronal environment. SGCs envelop neuronal cell bodies, whereas Schwann cells wrap the peripheral axons[9]. While Schwann cells are important for myelination, both SGCs and Schwann cells have well-documented roles in trophic and metabolic support of neurons and regeneration[10-13]. Immune cells, such as T cells and especially macrophages (including perineural macrophages), are prevalent in the human SG and DRG[14, 15]. In both ganglia, immune cells contribute to the neuro-immune interactions that modulate neuronal functions and responses to injury[16-21]. Various neuro-immune interactions in the DRG have been attributed to macrophages, which either promote or mitigate inflammation. Other non-neuronal cells, such as fibroblasts and vascular cells, have also been implicated in various painful conditions[22-24]. The intricate interactions and communications between these diverse non-neuronal cell types and the neurons are critical in both normal physiology and disease states, underscoring the importance of studying the cellular and molecular heterogeneity of the PNS in a high-resolution and cell-type-specific fashion.

Recent advancements in single-cell sequencing technologies have revolutionized our understanding of the diverse cell types present in the PNS [4, 15, 25-31], and their unique molecular characteristics underlying health and diseased conditions primarily in model animals [18, 19, 25, 28, 29, 32, 33]. However, understanding the molecular and cellular diversity of human PNS remains limited. In this study, we present an integrated single-cell multi-omic atlas of human thoracolumbar paravertebral SG and DRG using single-nucleus multiome sequencing (snMultiome-seq), which combines single-nucleus RNA sequencing (snRNA-seq) and single-nucleus assay for transposase-accessible chromatin using sequencing (snATAC-seq) from the same cells. By analyzing SG and DRG cells from multiple donors, we delineate the transcriptional and epigenomic landscapes of various cell types in these peripheral ganglia and their evolutionarily conserved and divergent features. We also identify key transcriptional similarities and differences in each cell type between SG and DRG, and the genomic regulatory networks that govern cell-type-specific gene expression in these peripheral ganglia. Our study provides an interactive companion website (https://peripheral-ganglia-multiome.shinyapps.io/atlas/) and offers new perspectives on the potential roles of diverse human PNS cell types in healthy biological systems and human diseased conditions broadly affecting various PNS tissues, such as hereditary sensory and autonomic neuropathies (HSAN).

## RESULTS

### High-quality human tissues enable multi-omic profiling of human peripheral ganglia at single-cell resolution

Technical advancements in recent years have transformed our understanding of the molecular and cellular diversity of the PNS in animal models, but multi-omic profiling of the human PNS has been hampered by unique challenges. Access to high-quality human tissues is limited, especially specimens with certain diseased and pathological states. Some existing tissues may not be ideal for single-cell multi-omic profiling, as progressive post-mortem RNA degradation and chromatin integrity loss can occur within a few hours and may be exacerbated by suboptimal extraction and storage conditions during the tissue procurement process[28, 34-37]. Moreover, profiling human PNS neurons is extremely difficult as they are proportionally much more sparse in human tissues compared to those in rodents and non-human primates[15, 28, 38]. Approaches, such as laser capture microscopy and spatial transcriptomics, offer superior neuronal coverage compared to traditional single-cell/single-nucleus sequencing techniques, but at the cost of under-representing non-neuronal cell populations[39-41]. Integrative, big data approaches may be necessary for an unbiased understanding of the cellular heterogeneity of human PNS[27, 42].

To address some of these challenges, we developed a human tissue procurement pipeline at Washington University in St. Louis, where we collaborate with MidAmerica Transplant to collect various post-mortem PNS tissues from organ donors[43]. In most cases, donor tissues are extracted and processed within one to three hours of cross-clamp. The high viability of the tissues enables both cell culture-based experiments[43-45] and molecular assays[15]. Frozen human PNS tissues derived from our tissue procurement pipeline exhibited excellent RNA (RNA integrity number [RIN]: 8.1 ± 0.4, n=9) and chromatin integrity (Figure S1A, n=8), allowing single-cell multi-omic profiling of the human PNS.

### A transcriptional and epigenomic atlas of human SG

As a single-cell multi-omic atlas of human SG is currently lacking, we first sought out to study the gene expression and chromatin accessibility of individual cell types present in human SG. We thus performed snMultiome-seq of post-mortem thoracolumbar paravertebral SG samples from seven individuals, generating eight snMultiome-seq libraries (Figure 1A, Table 1). After quality control processes and removal of low-quality and doublet nuclei (Methods), 52,634 nuclei were retained in the final dataset with an average of 30,697 paired reads sequenced and 1,810 genes detected per nucleus for the snRNA-seq libraries (Figure S1B), and 28,208 paired reads sequenced and 7,397 high-quality transposase-sensitive fragments for the snATAC-seq libraries. The transposase-sensitive fragments detected in the snATAC-seq libraries followed the expected nucleosomal size distribution (Figure S1A) and formed 139,026 accessible peaks when aggregated across all snATAC-seq libraries.

**Figure 1.**
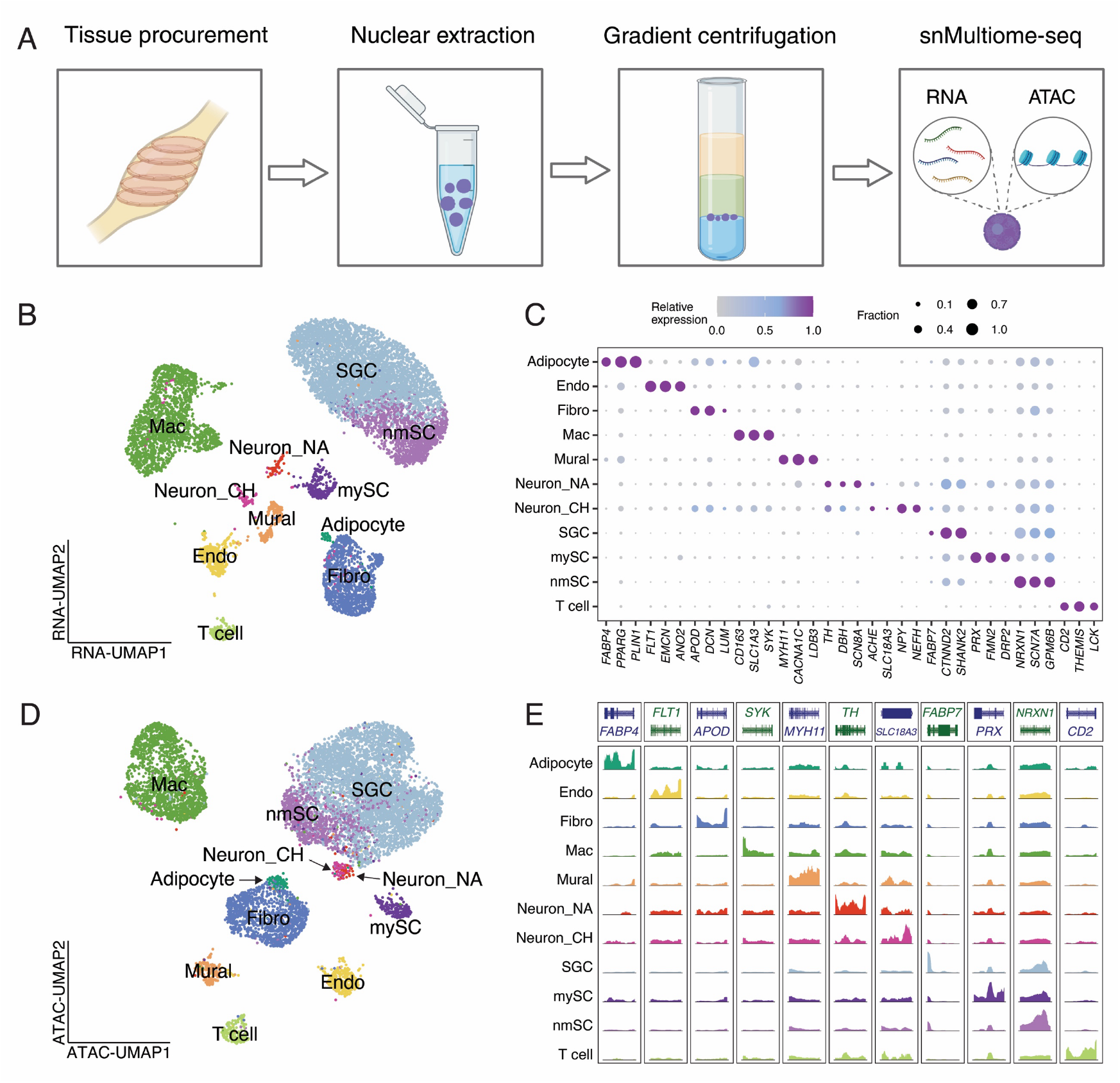
Transcriptional and epigenomic landscape of human SG. **A**. Experimental workflow of snMultiome-seq of human SG. **B**. Gene expression UMAP of human SG based on snRNA-seq data. 10,000 nuclei were randomly sampled and colored by transcriptional cell types. **C**. Dot plot displaying the expression of select marker genes in individual cell types. Dot size denotes the fraction of nuclei expressing a marker gene (>0 counts), and color denotes relative expression of a gene in each cell type (calculated as the mean expression of a gene relative to the highest mean expression of that gene across all cell types). **D**. Chromatin accessibility UMAP of human SG based on snATAC-seq data. 10,000 nuclei were randomly sampled and colored by transcriptional cell types. **E**. Coverage plot showing the accessibility around genomic loci of canonical cell-type-specific marker genes in each transcriptional cell type. The chromatin accessibility is displayed as the relative frequency of sequenced DNA fragments for each cell type, grouped by 50 bins per displayed genomic region. The frequency is normalized by the maximal frequency per genomic region. Endo: endothelial cell, Fibro: fibroblast, Mac: macrophage, SGC: satellite glial cell, mySC: myelinating Schwann cell, nmSC: non-myelinating Schwann cell. Neuron_NA: noradrenergic neuron, Neuron_CH: cholinergic neuron.

To identify transcriptionally distinct human SG cell types, we first performed dimension reduction and clustering of the snRNA-seq data. Notably, nuclei from individual libraries were represented in each cluster, indicating a minimal batch effect (Figure S1C-D). Clustering analysis revealed 11 major transcriptional cell types, including two neuronal types and nine non-neuronal cell types (Figure 1B,C), consistent with those previously described in human and mouse SG[4, 30, 40]. These cell types exhibit unique transcriptional profiles, and we identified 11,588 cell-type-specific marker genes (Log2FC > 1 and FDR < 0.05, Table 2) across all cell types. Among them, we found two neuronal types: noradrenergic neurons (Neuron_NA) and cholinergic neurons (Neuron_CH), which collectively represent 1.4% of the total population. Both neuronal types highly express genes important for synaptic neurotransmitter release, such as *SNAP25* (Synaptosome Associated Protein 25) and *SYT1* (Synaptotagmin 1, Figure S1E), and gene ontology (GO) analysis showed that their marker genes are associated with biological processes related to key neuronal functions, such as regulation of membrane potential and synaptic transmission (Figure S1G). The noradrenergic neurons express a high level of *TH* (Tyrosine Hydroxylase) and *DBH* (Dopamine Beta-Hydroxylase), both of which are essential for norepinephrine production (Figure 1C). The cholinergic neurons express *ACHE* (Acetylcholinesterase) and *SLC18A3* (encoding the vesicular acetylcholine transporter), genes crucial for cholinergic neurotransmission (Figure 1C). Cholinergic SG neurons may also be noradrenergic, as 61.3% and 57.5% of nuclei in the cholinergic cluster co-express *TH* and *DBH*, respectively (Figure S1F), although at a lower level compared to those in the nonandrogenic cluster. Notably, cholinergic neurons also express *NEFH*, encoding neurofilament heavy chain, at a higher level than noradrenergic neurons, suggesting their axons may be myelinated[46]. Our identification of the transcriptionally distinct noradrenergic and cholinergic SG neurons is largely consistent with those previously reported in humans and mice[4, 30, 40].

Glial cells are the most prevalent (60.5% of non-neuronal nuclei) non-neuronal cell types present in the human SG. We identified three glial cell types, including SGCs, myelinating Schwann cells (mySCs), and non-myelinating Schwann cells (nmSCs, Figure 1B-C). The SGCs distinctly express *FABP7* (Fatty Acid Binding Protein 7), a SGC marker gene previously validated in peripheral ganglia across species[4, 15]. mySCs express *PRX* (Periaxin), an essential gene for the maintenance of peripheral nerve myelin, and nmSCs express *NRXN1* encoding Neurexin 1, a nmSC marker encoding a cell adhesion molecule important for the formation and function of synapses [4, 15]. These glial cell types have distinct roles in supporting, protecting, and regulating the microenvironment of PNS neurons[10, 13, 47, 48]. Consistent with this notion, we found glial cell marker genes are associated with GO terms, such as synaptic regulation, ion transmembrane transport, axon guidance and myelination, as well as cell adhesion (Figure S1G). Moreover, we also identified endothelial cells expressing *FLT1*, fibroblasts expressing *APOD*, mural cells expressing *MYH11*, adipocytes expressing *FABP4*, as well as immune cells, such as *SYK*-expressing macrophages and *CD2*-expressing T cells (Figure 1B-C). The marker genes expressed in these cell types are also associated with their unique functions in the human SG (Figure S1G). All cell types are present in each library except for adipocytes derived from a subset of donors (Figure S1D), which are likely due to the accidental inclusion of surrounding tissues during the tissue procurement and processing.

To study the chromatin accessibility profiles of human SG cell types, we performed dimension reduction and clustering of the snATAC-seq data generated from the same nuclei assayed in snRNA-seq, generating 26 epigenomically distinct clusters (Figure S2A). While most cell types were well represented on the UMAP, several SGC clusters appeared to exhibit donor-biased representation (Figure S2A), likely reflecting epigenomic differences underlying the donors’ history, or introduced by technical variations during tissue procurement and sequencing process. Expectedly, 46.2% of all snATAC-seq peaks were donor-specific (accessible in less than half of the donors, Figure S2C). To control for donor variability, we next excluded these donor-specific peaks from the chromatin accessibility analysis, resulting in an improved clustering with reduced bias against libraries and donors (Figure S2B). The post-correction chromatin accessibility exhibited a high level of correlation with the gene expression, as 88.4 ± 12.3% of nuclei in each snATAC-seq cluster are from the same transcriptional cell types previously assigned (Figure S2D). To harmonize the cell type annotation across multi-omic analyses, we subsequently grouped nuclei by transcriptional cell types for all downstream analyses. The cell types display unique epigenomic profiles (Figure 1D) and distinct chromatin accessibility around the genomic loci of canonical marker genes (Figure 1E). We identified 43,496 cell-type-specific peaks (Log2FC > 1, FDR < 0.05, Table 2) across all human SG cell types.

### Putative sex difference in SG cell types

Several sex-specific differences in human autonomic functions have been previously described[49-51], which prompted us to investigate whether these differences may be driven by unique transcriptional and epigenomic features in the SG samples between three males and two females that were sequenced individually. Overall, the male and female libraries exhibit similar cell type distribution (Figure S3A). Male and female SG also display similar transcriptional and epigenomic landscapes (Figure S3B,C), with strong correlations in gene expression (Pearson’s *r* = 0.95 ± 0.05 per cell type) and chromatin accessibility (*r* = 0.74 ± 0.18) in individual cell types.

Despite the similarity, differential expression analysis comparing nuclei of the same cell type between the sexes revealed 82 genes significantly enriched (Log2FC > 1, FDR < 0.05) in males and 50 genes enriched in females (Figure S3D, Table 3). The most common sex differences across cell types are known sex-specific genes involved in X inactivation (such as *XIST* and *TSIX*) or are Y chromosome genes (such as *KDM5D, UTY*, and *DDX3Y*). Consistently, differential chromatin accessibility is also observed in genomic loci of these known sex-specific genes between nuclei from male and female donors (Figure S3E). In addition, we identified differentially expressed genes between male and female cell types (Figure S3F, Table 3). For example, the expression of *NPY* (Neuropeptide Y), important for stress response, is 9.3-fold significantly higher in endothelial cells in females than males. The expression of *GRM7*, which encodes metabotropic glutamate receptor 7, is 6.7-fold higher in in cholinergic neurons from males than females (Figure S3F). *GRM7* is broadly expressed in the CNS and is important for modulating neurotransmitter release, but its sex-specific expression and potential functions in the PNS have not been reported. In addition to sex-specific gene expression, we identified 278 male-specific peaks and 75 female-specific peaks (Log2FC > 1, FDR < 0.05) in snATAC-seq data that may underlie sex-specific transcription regulation (Table 3). These putative transcriptional and epigenomic sex-specific features may be important for the sex-specific SG functions. Given the small donor numbers for each sex in our snMultiome-seq data, future studies will be required to validate these putative sex-specific features.

### Transcriptional conservation between human and mouse SG

Understanding the molecular and cellular evolutionary convergence and divergence of SG is crucial, as the species-specific features may contribute to species-specific functions, potentially limiting the predictive value of mouse models for understanding human biology. Despite recent advances in single-cell transcriptomics that have enabled the molecular characterization of mouse SG cell types, it remains unclear to what extent the SG cell types and their cell-type-specific molecules are evolutionarily conserved. To directly address these questions, we compared our snRNA-seq data generated from human thoracolumbar paravertebral SG to that derived from mouse superior cervical SG previously reported[4]. When performing clustering without integration, the data are separated by species (Figure S4A), likely due to a combination of species-specific, spinal-level differences, and technical variations (e.g., different sample collection procedures and sequencing techniques). To better compare the cell types present in humans and mice, we integrated the two datasets based on their shared variable features[52]. After integration, nuclei of the same cell types from the two species were clustered together (Figure 2A), resulting in highly similar cell type correspondence (88.6% ± 12.8% per cell type, Figure 2B).

**Figure 2.**
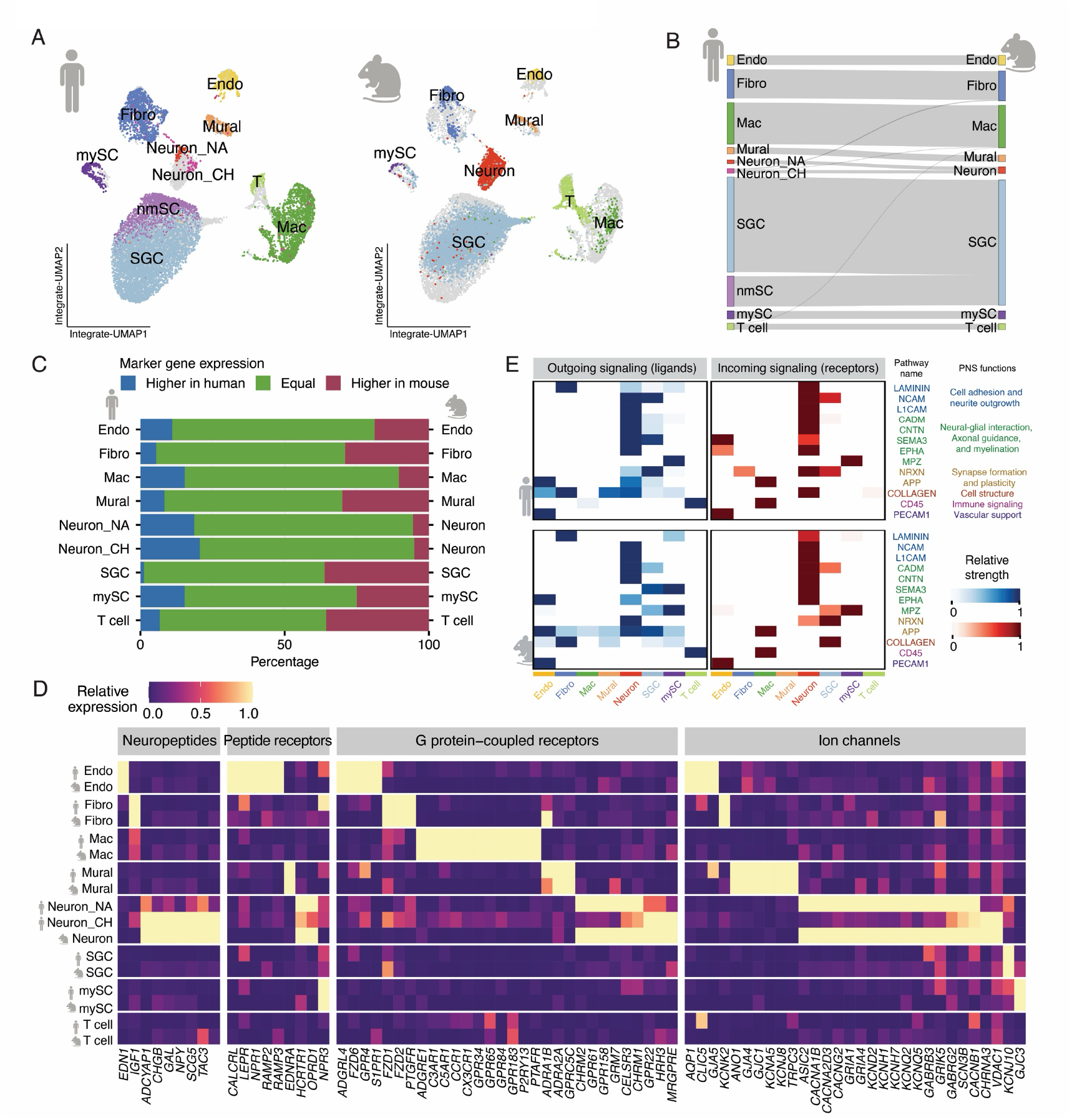
Cross-species conservation of SG cell types. **A**. Integrative gene expression UMAP of the human and mouse SG based on snRNA-seq data. 10,000 nuclei were randomly sampled per dataset for visualization. Left: human nuclei are colored by cell types, and mouse nuclei are displayed in the background as grey dots. Right: mouse nuclei are colored by cell types previously reported, and human nuclei are displayed in the background as grey dots. **B**. Sankey plot showing the similarities between human and mouse SG cell types. Gray lines connect the human cell types (left) to the previously reported mouse classifications (right), indicating the cell types to which they are most closely related, as quantified by the co-clustering of human nuclei in each cell type with the respective mouse cell types when the two datasets were integrated. **C**. Bar plot showing the percentage of cell-type-specific marker genes for each cell type grouped by their expression level in human and mouse. Pairwise differential expression analysis was performed comparing the nuclei of the paired cell types between humans and mice. Genes with Log2FC >1 and FDR < 0.05 are higher in human, genes with Log2FC < (−1) and FDR < 0.05 are higher in mouse, and all other genes are equally expressed in the two species. **D**. Heatmap displaying the expression of select cell-type-specific physiologically relevant genes in individual human and mouse SG cell types. Color denotes relative expression of a gene in each cell type of each species (calculated as the mean expression of a gene relative to the highest mean expression of that gene across all cell types of a given species). **E**. Heatmap showing the relative strength of conserved signaling pathways across individual human (top panels) and mouse (bottom panels) cell types. Cell types not present in both species were excluded, and human neuronal clusters were merged as one neuronal cell type for ligand receptor analysis. Endo: endothelial cell, Fibro: fibroblast, Mac: macrophage, SGC: satellite glial cell, mySC: myelinating Schwann cell, nmSC: non-myelinating Schwann cell. Neuron_NA: noradrenergic neuron, Neuron_CH: cholinergic neuron.

To gain insight into the evolutionary conservation of SG at the molecular level, we next compared the cell-type-specific transcriptional profiles in individual SG cell types between humans and mice. The overall transcriptional profile in individual cell types is largely similar, as the expression levels of cell-type-specific marker genes are highly correlated between human and mouse SG (*r* = 0.79 ± 0.08 per cell type). In addition, 57.7 – 75.8% of the cell-type-specific marker genes had similar expression levels between human and mouse cell types (less than two-fold difference of expression between the species, Figure 2C). We also observed very similar gene expression patterns for some physiologically relevant genes encoding key neuropeptides and receptors, G-protein coupled receptors (GPCRs), and ion channels between human and mouse SG cell types (Figure 2D).

Moreover, to compare the cell-cell communications in human and mouse SG, we performed ligand-receptor analysis on each dataset separately. To control the effect of technical variation, we downsampled the human data to match the mouse data (Methods). In total, we identified 28 and 26 significantly enriched signaling pathways in human and mouse SG, respectively. Among them, 13 signaling pathways are shared between species (Figure 2E). These evolutionarily conserved pathways are associated with important SG cell functions such as neurite outgrowth, myelination, and synaptic functions, and several genes have been implicated in various human diseases. For example, APP (amyloid precursor protein) and NCAM (Neural cell adhesion molecule) signaling pathways are crucial for neuronal development and synapse formation in the nervous system[53-55]. In both human and mouse SG, *APP* is expressed in neurons and several non-neuronal cell types, such as SGCs and fibroblasts (Figure 2E, S5A,E,F). NCAM ligands *NCAM1, NCAM2*, and the receptor *L1CAM* are expressed in SG neurons in both species, whereas *NCAM1* and *NCAM2* are expressed in SGCs and Schwann cells in humans but not in mice (Figure 2E, S5B,E,F). CNTN (Contactin) and LAMININ play physiological roles in axonal and cellular structural support, and *CNTN1* and *LAMA2* are expressed in various SG cell types in humans and mice (Figure 2E, S5C-F). Abnormality of APP and NCAM in the SG has been implicated in animal models of neurodegenerative disorders, such as dementia[56], and *CNTN1* and *LAMA2* genes are directly link to certain types of congenital muscular dystrophies in human populations[57, 58]. Since little is known about the mechanisms by which these molecular perturbation contribute to the autonomic dysfunctions commonly observed in patient populations, this analysis explores the potential molecular and cellular basis of SG that may be affected and conversely contribute to the autonomic dysregulations reported in human diseases[59, 60]. This analysis also sheds light on the evolutionary conservation that would aid the future investigation of human diseases using animal models.

### Cellular and molecular divergence of human and mouse SG

Despite the overall molecular and cellular conservation between human and mouse SG, we also observed some differences between the two datasets that may suggest species-specific SG functions. On the cellular level, neuronal populations in mice are 14.2 times more abundant than those in humans (Figure S4B). The lower relative abundance of neurons versus non-neuronal cells in human SG is consistent with previous studies of other peripheral ganglia[15, 28]. In contrast to the neuronal populations identified in human data, all neuronal clusters in the mouse data appear to be noradrenergic with a lack of cholinergic marker expression[4]. Notably, mouse neurons are transcriptionally more similar to human noradrenergic neurons compared to human cholinergic neurons (Figure S4C). This observation is consistent with previous reports that cholinergic neurons are more abundant in human SG than mouse SG[40, 61, 62]. mySCs, which support myelination, are also more abundant in human data than in mouse data (Figure S4B). We also observed proportionally more abundant macrophages in humans than in mice (Figure S4B), which is consistent with previous cross-species comparison of peripheral ganglia and may contribute to species-specific neuro-immune interactions[27, 28].

In addition, we found 1,010 genes differentially expressed between human and mouse SG cell types, including 18 physiologically relevant genes more highly expressed in human cell types, and 30 physiologically relevant genes more highly expressed in mouse cell types (Figure S6A). Moreover, we identified 15 signaling pathways that are significantly enriched in the human data and 13 signaling pathways significantly enriched only in the mouse data (Figure S6B,C). Among the species-specific signaling pathways, glutamate signaling is enriched only in the human data (Figure S6D). *SLC1A3* (encoding the Excitatory Amino Acid Transporter 1) is distinctly expressed in human macrophages. *GRIK2* and *GRIK3*, which encode the Glutamate Ionotropic Receptor Kainate Subunits 2 and 3, respectively, are expressed in human SG neurons and non-neuronal cells, such as endothelial cells, fibroblasts, SGCs, and Schwann cells. However, *Slc1a3, Grik2*, and *Grik3* are not significantly detected (< 5% nuclei per cell type) in any mouse SG cell types. Serotonin/dopamine signaling is enriched only in the mouse data (Figure S5E). *Ddc* (Dopa Decarboxylase) and *Htr3a* (encoding 5-Hydroxytryptamine [serotonin] Receptor 3A) are highly expressed in mouse SG neurons, while not significantly detected in any human SG cell types. Some of these differences between the human and mouse SG data may be important for species-specific SG signaling. Future studies are required to validate the species-specific differences reported in this study.

### Transcriptional comparison between human SG and DRG

The sensory system in the DRG relays ascending signals from the environmental stimuli and internal organs to the CNS, while the sympathetic system relays descending signals from the CNS to the peripheral tissues for mediating the ‘‘fight or flight’’ response and maintaining body homeostasis. Despite the shared developmental origin, DRG and SG exhibit distinct innervation patterns and carry out unique physiological roles and functions. Previous studies reported that SG and DRG share similar cell types in rodents[4], but the extent of transcriptional and epigenomic similarities and differences in the human SG and DRG cell types has not been examined.

To address these questions, we integrated our snMultiome-seq data from human SG with the snMultiome-seq data from human DRG reported in a companion study[38]. As the tissues used in both studies were derived from the same procurement pipeline and processed and sequenced using the same experimental protocols, this allowed us to rigorously control for technical variations. Indeed, we identified overlapping cell types from the two datasets with similar cell type distribution (Figure S7A). Human SG and DRG cell types exhibit high levels of transcriptional similarity, as nuclei of the same cell types from SG and DRG co-clustered nicely on the UMAP, even before integration (Figure 3A, S7B,C), and 74.7 – 99.7% of nuclei from each SG cell type are anchored to the same DRG cell type in the integration analysis (Figure 3B). When we examined the cell-type-specific transcriptional profiles, 82.8 – 98% of the cell-type-specific marker genes in each cell type are expressed at a similar level (less than two-fold difference between the two ganglia (Figure 3C). Interestingly, the human SG and DRG neuronal subtypes exhibited a high level of transcriptional correspondence. 80.1% of SG noradrenergic neurons are mapped to DRG neuron_1 cluster (Figure 3B), which transcriptionally matches the unmyelinated small-diameter C-fiber neuronal populations in the DRG[38]. SG noradrenergic neurons and DRG neuron_1 cluster are transcriptionally more similar to each other than to other SG and DRG neuronal types (Figure S7D). Out of 6,661 marker genes that are expressed in these two DRG and SG neuronal types, 85.3% are expressed at similar levels. These two neuronal types both distinctly express ion channels and GPCRs, such as *SCN9A, OPRD1*, and *OPRM1* (Figure S7E). Similarly, 99.7% of SG cholinergic neurons are mapped to DRG neuron_2 cluster (Figure S7D), which transcriptionally aligns with the myelinated large-diameter A-fiber DRG neuronal populations[38]. They are transcriptionally more similar to each other than to other SG and DRG neuronal types, and 87.5% of the 3,314 marker genes are expressed at similar levels. SG cholinergic neurons and DRG neuron_2 both distinctly express genes associated with myelinated axons, such as *NEFL, NEFH*, and *NEFM*.

**Figure 3.**
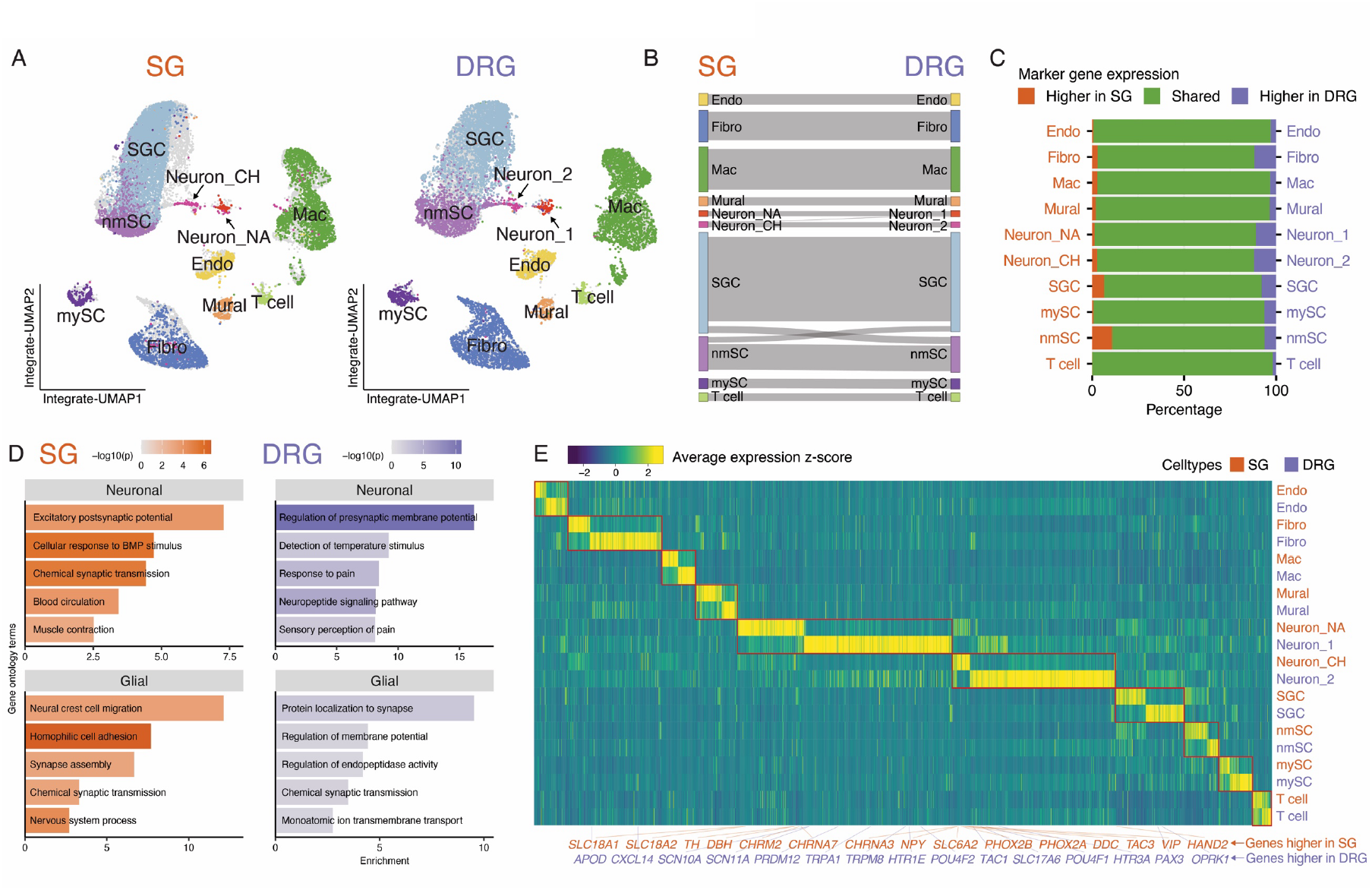
Comparison between human SG and DRG. **A**. Integrative gene expression UMAP of the human SG and DRG based on snRNA-seq data. 10,000 nuclei were randomly sampled per dataset. Left: SG nuclei are colored by cell types, and DRG nuclei are displayed in the background as grey dots. Right: DRG nuclei are colored by cell types previously reported, and SG nuclei are displayed in the background as grey dots. **B**. Sankey plot showing the similarities between human SG and DRG cell types. Gray lines connect the SG cell types (left) to the previously reported DRG cell type classifications (right) they are most closely related, quantified as the SG nuclei in each cell type co-clustered to the respective DRG cell types when the two datasets were integrated. **C**. Bar plot showing the percentage of cell-type-specific marker genes for each cell type grouped by their expression level in SG and DRG. Pairwise differential expression analysis was done comparing nuclei of the same cell type between SG and DRG. Genes with Log2FC >1 and FDR < 0.05 are higher in SG, genes with Log2FC < (−1) and FDR < 0.05 are higher in DRG, and all other genes are equally expressed in the two ganglia. **D**. Plot showing the top five biological processes enriched in the GO analysis of SG and DRG. **E**. Heatmap displaying the expression of cell-type-specific genes that are differentially expressed between SG and DRG (See Table 4 for the original data). Cell types not present in both ganglia were excluded. SG: sympathetic ganglia, DRG: dorsal root ganglia. Endo: endothelial cell, Fibro: fibroblast, Mac: macrophage, SGC: satellite glial cell, mySC: myelinating Schwann cell, nmSC: non-myelinating Schwann cell. Neuron_NA: noradrenergic neuron, Neuron_CH: cholinergic neuron. Neuron_1: largely C-fiber neuron, Neuron_2: largely A-fiber neuron.

Despite the overall transcriptional similarity, SG and DRG cell types also exhibit some key differences that may underlie their distinct neurobiological functions. In total, we identified 348 genes more highly expressed in SG cell types (Log2FC>1, FDR<0.05) and 769 genes more highly expressed in DRG cell types (Table 4). SG and DRG neurons differ in many key aspects, such as morphology, neurotransmitter release, physiology, and function, which appeared to be reflected in their gene expression patterns. SG-enriched genes are important for autonomic regulatory function, such as regulation of vasculature, muscle contraction, and blood circulation (Figure 3D). Compared to DRG, SG neurons more highly express genes important for norepinephrine production and signaling (e.g., *SLC18A1, SLC18A2, TH, DBH, SLC6A2, and DDC*), and cholinergic signaling (e.g., *CHRNA3* and *CHRNA8* encoding subunits of the nicotinic acetylcholine receptor genes, and *CHRM2* encoding a muscarinic acetylcholine receptor, Figure 3E). SG neurons also more highly express neuropeptide genes, including *NPY* (Neuropeptide Y), *VIP* (Vasoactive intestinal peptide), and *TAC3* (encoding Neurokinin B). Genes encoding transcription factors crucial for the sympathetic cell type development and specificity, such as *PHOX2A, PHOX2B*, and *HAND2*, are also more highly expressed in human SG than DRG. On the other hand, DRG-enriched genes are important for the detection of environmental stimuli and sensory perceptions, including pain (Figure 3D). Compared to SG, DRG neurons highly express genes encoding ion channels for nociceptive signal transmission (e.g., *SCN10A* and *SCN11A*), temperature sensing (*TRPA1* and *TRPM8*, Figure 3E). DRG neurons also more highly express *SLC17A6*, encoding a vesicular transporter of glutamate, the primary neurotransmitter released by the DRG neurons, and *TAC1*, encoding substance P, a key modulator for pain sensation[63, 64]. Neuropeptide receptor genes, such as serotonin receptors (*HTR1E* and *HTR3A*) and opioid receptor *OPRK1*, are more highly expressed in DRG than SG. Sensory neuron-specific transcription factor genes, such as *PRDM12, PAX3, POU4F1*, and *POU4F2*, are also more highly expressed in DRG than SG.

### Glial cell type distribution is similar in human SG and DRG

SGCs and Schwann cells are the main types of glial cells in the PNS. SGCs envelop the cell bodies of neurons, whereas Schwann cells ensheathe axons[10, 11, 65]. A previous comparative study has shown that mySCs are more sparse in the mouse superior cervical SG compared to the mouse lumbar DRG, whereas the SGCs are similarly abundant [4]. However, both SGCs and mySCs are similarly abundant in our human SG and DRG atlas. To validate our findings, we investigated the distribution of glial cell types in human SG and DRG using RNA fluorescent in-situ hybridization (RNA-FISH, Figure 3). We labeled mySCs with *PRX*, which is specifically expressed in mySCs (Figure 4G). In addition, we probed for *NRXN1* to label SGCs and nmSCs. While *NRXN1* is broadly expressed in all cell types (Figure S8A), its expression level is 5.7-fold higher in nmSCs and 1.4-fold higher in SGCs than in other cell types (Figure 4G). Therefore, we reasoned that *NRXN1* can preferentially label nmSCs and SGCs based on different expression levels. Finally, we used *SNAP25*, a pan-neuronal marker, to label the neuronal population.

**Figure 4.**
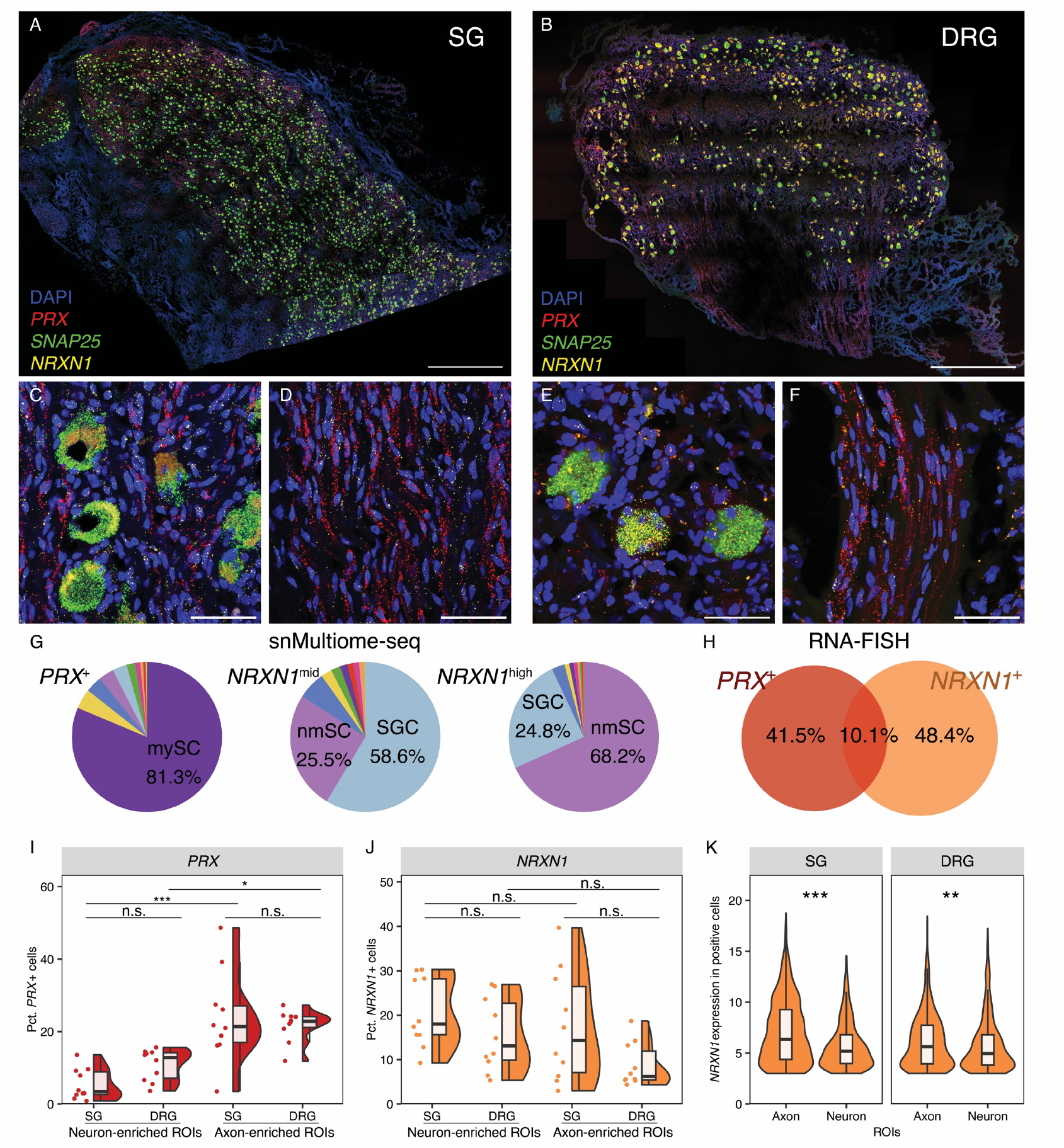
Glial cell distribution between human SG and DRG. **A**. A representative FISH image from human SG. Scale bar = 1mm. **B**. A representative FISH image from human DRG. Scale bar = 1mm. **C and E**. Representative zoomed-in images of neuron-enriched regions in human SG (C) and DRG (E). Scale bar = 50 μm. **D and F**. Representative zoomed-in images of axon-enriched regions in human SG (D) and DRG (F). Scale bar = 50 μm. **G**. Cell type distribution of *PRX*+ (> the median value among all positive nuclei) nuclei and *NRXN1* mid- (> the median value and no more than 75-percentile among all positive nuclei) and high-expressing (> 75-percentile among all positive nuclei) nuclei in snMultiome-seq data of human SG and DRG. Color denotes cell types. **H**. The overlap between *PRX*+ cells and *NRXN1*+ cells in RNA-FISH analysis. **I**. Quantification of *PRX*+ cell distribution across different ROIs. Two-way ANOVA: F (3,36) = 14.32, p = 2.6E-6 [***]. Tukey HSD tests: p = 0.38 (neuron DRG-neuron SG), p = 1.3E-5 [***] (axon SG-neuron SG), p = 0.57(axon DRG-axon SG), p = 0.014 [*] (axon DRG-neuron DRG). **J**. Quantification of *NRXN1*+ cell distribution across different ROIs. Two-way ANOVA: F (3,36) = 3.26, *p* = 0.03 [*]. Tukey HSD tests: *p* = 0.38 (neuron DRG-neuron SG), *p* = 0.83 (axon SG-neuron SG), *p* = 0.15(axon DRG-axon SG), *p* = 0.32 (axon DRG-neuron DRG). **K**. The distribution of *NRXN1* expression from different ROIs in human SG and DRG. One-way t-test testing if expression is higher in axon-enriched ROIs than neuron-enriched ROIs: *p* = 2.1E-28 [***] for SG, and p = 0.0015 [**] for DRG. [\**p* = 0.05, \*\*\**p* = 0.001]. SG: sympathetic ganglia, DRG: dorsal root ganglia. SGC: satellite glial cell, mySC: myelinating Schwann cell, nmSC: non-myelinating schwann cell.

We performed RNA-FISH on thoracolumbar paravertebral SG and DRG samples from two donors (Table 1, Figure 4A,B). For each sample, we analyzed five neuron-enriched regions of interest (ROIs) and five axon-enriched ROIs (Figure 4C-F). We then segmented the cells using cellpose and quantified the expression and called positive cells using SCAMPR (Figure S8B-D, Methods)[66, 67]. In total, we identified 38,034 SG cells and 50,426 DRG cells. *PRX* labeled 13,132 (14.8%) cells and *NRXN1* labeled 14,901 (16.8%) cells. *PRX* and *NRXN1* labeled distinct populations, both of which showed little expression of *SNAP25* (Figure 4H). Consistent with our human sequencing data, we found that both *PRX*+ cells and *NRXN1*+ cells are distributed similarly between human SG and DRG (Figure 1I,J). Our snMultiome-seq and RNA-FISH results collectively show that individual glial cell types are similarly abundant between human SG and DRG, in contrast to the mouse, where mySCs are less abundant in the SG than DRG. This may suggest a potential cross-species difference.

Expectedly, in both SG and DRG, *PRX*+ cells are more enriched around axon-enriched regions compared to the neuron-enriched regions, suggesting that mySCs are preferentially localized around axonal bundles. While *NRXN1*+ cells are relatively equally distributed across different regions in SG and DRG, *NRXN1* expression is higher in the axon-enriched regions than neuron-enriched regions (Figure 4K). These results suggest that nmSCs may be more enriched around axons, and *NRXN1* lower-expressing SGCs may be more enriched around neuronal somas.

### Genomic regulatory networks of human peripheral ganglia

While the gene expression programs in the diverse PNS cell types have been largely elucidated in animals and in some human tissues, the genomic regulatory networks (GRNs) that govern the distinct gene expression patterns in individual cell types have not yet been explored in humans. The intricate interactions between the trans-acting genomic regulatory elements (GRE, e.g., transcription factors [TF]) and cis-acting GREs (e.g., promoters, enhancers, and silencers) fine-tune the gene expression in a cell-type-specific manner. Our multi-omic atlas of the human peripheral ganglia offers a valuable resource to characterize these putative GREs that control the cell-type-specific gene expression and study the GRNs underlying the distinct identities and functions of individual human PNS cell types. Our study might also provide new opportunities for understanding the function of disease-associated genetic variations and designing cell-type-specific genetic tools (e.g., transgenic mice or gene therapy vectors).

To study the cell-type-specific GRNs, we performed Functional Inference of Gene Regulation (FigR) analysis on the snMultiome-seq data of human SG and DRG[68]. By correlating the accessibility of snATAC-seq peaks with the expression of cell-type-specific genes in snRNA-seq, we were able to connect the distal GREs to target genes and infer the activity of TFs that may mediate these regulatory events. In total, we identified 39 TFs whose regulatory activities are enriched in human SG and/or DRG (Figure 5A), Among them, 25 TFs are shared between the two ganglia, including 15 transcription activators, whose motif enrichment is positively correlated with the expression of regulated genes, and 10 transcription repressors, with negative correlation between motif enrichment and gene expression (Figure 5B). For example, SOX10 (SRY-Box Transcription Factor 10) is a key regulator of neural crest cell development and critical for the development of glial cells across different PNS tissues[69-71]. Indeed, we found that the SOX10 gene is highly expressed in the SGCs, mySCs, and nmSCs in both human SG and DRG, and the SOX10 binding motif is also highly enriched in the accessible peaks of these cell types (Figure 5C). We identified 354 genes putatively regulated by SOX10 (Figure 5D), which significantly overlap with cell-type-specific marker genes for glial cell types (Figure S9A, Table 5). These results suggest that SOX10 may act as key TF for glial cell types in both human SG and DRG by activating and maintaining the expression of marker genes for different glial cell types. Similarly, SPI1 (Spi-1 Proto-Oncogene, also known as PU.1) is well known for its role in the development and differentiation of immune cells[72-74]. SPI1 is likely a transcription activator for the SG and DRG macrophages due to its high gene expression and motif enrichment (Figure 5E). The majority of the SPI1-regulated genes is distinctly expressed in macrophages (Figure 5F), and SPI1 may be a key regulator for macrophage functions as its top regulated genes are associated with immune processes such as inflammatory response (*SYK, ALOX5, C5AR1, CD14, TAB2*), immune signaling (*THEMIS2, MS4A7, FPR3*), and pathogen recognition and processing (*NCF2, CTSB*, Figure 5G, Table 5).

**Figure 5.**
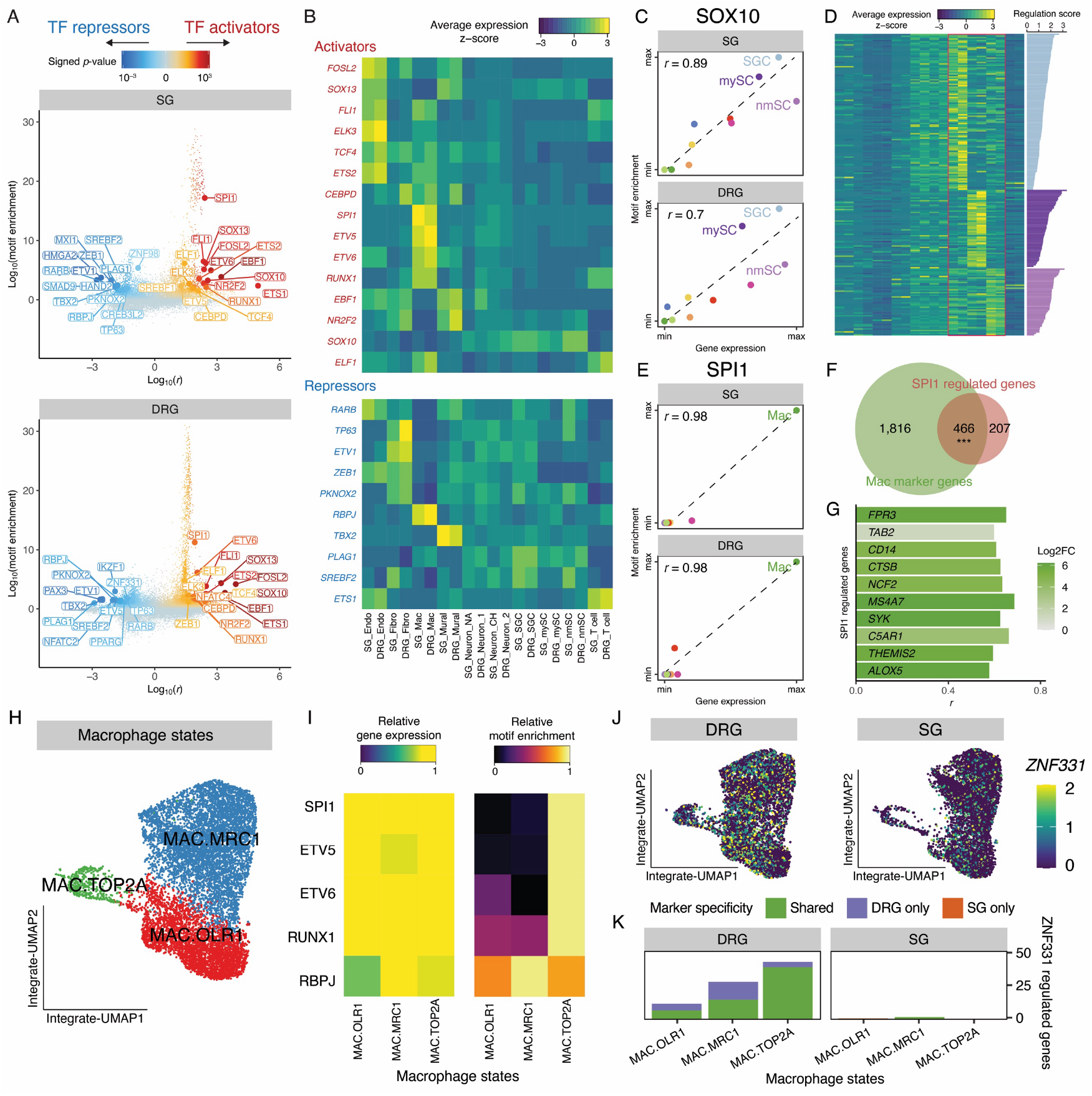
Genomic regulatory networks of human peripheral ganglia. **A**. Scatter plot showing the TFs enriched in human SG (top) and human DRG. Each dot shows the interaction between one TF and one of its regulated genes. For each TF, the interaction with the highest score is highlighted and labeled. **B**. Heatmap showing the expression of TF genes in individual SG and DRG cell types based on the snRNA-seq data. **C**. Scatter plot of the gene expression and motif enrichment of SOX10 in individual SG (top) and DRG (bottom) cell types. Pearson’s r and the line of best fit are displayed. **D**. Heatmap showing the expression of SOX10-regulated marker genes in satellite glial and Schwann cells. The row of the heatmap follows the same order as Figure 5B. Bars show the regulation scores of individual genes by SOX10. **E**. Scatter plot of the gene expression and motif enrichment of SPI1 in individual SG (top) and DRG (bottom) cell types. Pearson’s r and the line of best fit are displayed. **F**. Overlap between SPI1-regulated genes and macrophage-specific marker genes. Hypergeometric test: p = 6.8E-82 [***]. **G**. Bar plot showing the top 10 SPI1-regulated genes based on regulation score. Colors indicate the differential expression of genes in macrophages compared to all other cell types. **H**. Integrative gene expression UMAP showing the subtypes of 10,000 randomly sampled macrophage nuclei. **I**. Heatmaps showing the gene expression (left) and motif enrichment (right) of TFs in macrophage subtypes. **J**. Integrative gene expression UMAP showing the expression of *ZNF331* in 5,000 SG (right) and 5,000 DRG (left) macrophage nuclei randomly sampled. **K**. Number of genes predicted to be regulated by ZNF331 in individual DRG and SG macrophage subtypes. SG: sympathetic ganglia, DRG: dorsal root ganglia. Endo: endothelial cell, Fibro: fibroblast, Mac: macrophage, SGC: satellite glial cell, mySC: myelinating Schwann cell, nmSC: non-myelinating Schwann cell. Neuron_NA: noradrenergic neuron, Neuron_CH: cholinergic neuron. Neuron_1: largely C-fiber neuron, Neuron_2: largely A-fiber neuron.

Besides the shared TFs between the two peripheral ganglia, we also identified eight TFs enriched only in SG and six TFs enriched only in DRG (Figure S9B,D), including HAND2 (Heart And Neural Crest Derivatives Expressed 2), important for the development of the autonomic nervous system including the noradrenergic neurons[75, 76]. Its gene expression and motif enrichment are only detected in SG (Figure S9C). PAX3 is pivotal for the differentiation of neural crest precursor cells into sensory neurons[77, 78]. Similarly, its gene expression and motif enrichment are only detected in DRG (Figure S9E). These TFs may be important for the development and differentiation of the human SG and DRG, underscoring their unique properties and functions.

Interactions between neurons and the immune system are important for regulating neuronal functions and have been implicated in numerous pathophysiological conditions, such as obesity and chronic pain[19, 79-81]. Macrophages in the DRG are a cell type of interest since they have roles in regulating neuropathic pain, axonal regeneration, and the ganglia microenvironment in animal studies[21, 82], and there is a population of macrophage/macrophage-like cells that is in close proximity to neurons after injury in both animal models and in humans[14, 83, 84]. In our analysis, we identified six TFs that exhibited the highest expression in macrophages, including those that are shared between SG and DRG (SPI1, ETV5, ETV6, RUNX1, and RBPJ), and ZNF331 exhibiting much higher activity in DRG than SG. To further investigate the functions of these macrophage-specific TFs, we performed subclustering of the macrophages and identified three clusters with relatively equal representation from both SG and DRG nuclei (Figure 5H, S10A). These clusters likely represent different macrophage states based on their marker genes: MAC.OLR1 expresses pro-inflammatory markers, such as *OLR1* and *CD83*, whereas MAC.MRC1 expresses anti-inflammatory markers, such as *CD163* and *MRC1* (Figure S10B). MAC.TOP2A expresses markers of proliferation, including *TOP2A, MKI67*, and *BUB1B*. These macrophage clusters exhibit distinct gene expression patterns with enriched GO terms suggesting their potential functions (Figure S10C, Table 6). To better understand the molecular mechanisms that may govern the diverse macrophage functions, we next investigated the TFs and their GRNs in different macrophage clusters. While the expression of SPI1, ETV5, ETV6, and RUNX1 is similar across macrophage subtypes in snRNA-seq data, their motifs are most enriched in MAC.TOP2A (Figure 5I), suggesting that, in addition to their roles in macrophage differentiation, those TFs may be especially important for macrophage proliferation and maintenance in human SG and DRG. While not experimentally confirmed, these data support the hypothesis that, similar to the mouse, there is a self-renewing population of macrophages in human peripheral ganglia[83, 84]. The RBPJ gene is expressed 2.7-fold higher in anti-inflammatory MAC.MRC1 than in other subtypes, whereas its motif enrichment is also higher in MAC. MRC1 (Figure 5I). RBPJ may act as a transcription activator for controlling macrophage gene expression and polarization of anti-inflammatory states. Moreover, *ZNF331* is expressed 1.5 – 2.1 times higher in different macrophage clusters of human DRG than SG (Figure 5J), and predicted to regulate 83 genes in DRG macrophages, compared to only three genes in SG counterparts (Figure 5K, Table 6). Therefore, ZNF331 may be key to drive DRG macrophage gene expression signatures, and functional assessment of ZNF331 control of DRG, but not SG, macrophage inflammatory states will be an important future step.

### Cell types implicated in hereditary sensory and autonomic neuropathies

Single-cell atlases are useful for identifying the cell types and genes that may contribute to human diseases[28, 41, 85]. Hereditary sensory and autonomic neuropathies (HSAN) are a group of genetically and phenotypically heterogeneous rare disorders. To date, nine types of HSAN have been identified based on the age of onset, clinical features, and genetic inheritance (types I to IX, Figure 6A), with several types further divided into multiple subtypes[86-90]. The key clinical features of HSAN include sensory dysfunctions (insensitivity to pain and temperature, less affected touch sensation, and proprioception) and varying degrees of autonomic deficits (e.g., gastrointestinal dysfunction, cardiovascular instability, and anhidrosis or hypohidrosis), with some types also exhibiting motor symptoms[91, 92]. These clinical manifestations suggest that HSAN broadly affects the human PNS, but the effect may vary across sensory, autonomic, and motor components. It is thought that each HSAN-associated mutation causes disruption in different biological processes in the sensory and autonomic systems (and motor system in some types), but the neurobiological mechanisms by which the mutant genes affect normal functions of diverse human PNS cell types remain elusive. As a result, there is no treatment currently available to address the underlying cause and/or prevent disease progression for HSAN patients. As pathological human tissues from HSAN patients are extremely rare, we reasoned that our multi-omic atlas of the human peripheral ganglia could provide a new opportunity for localizing and predicting the cell types in which genomic variations drive HSAN pathophysiology by examining the expression of HSAN-associated genes and chromatin accessibility of HSAN-associated GREs in a cell-type-specific fashion.

**Figure 6.**
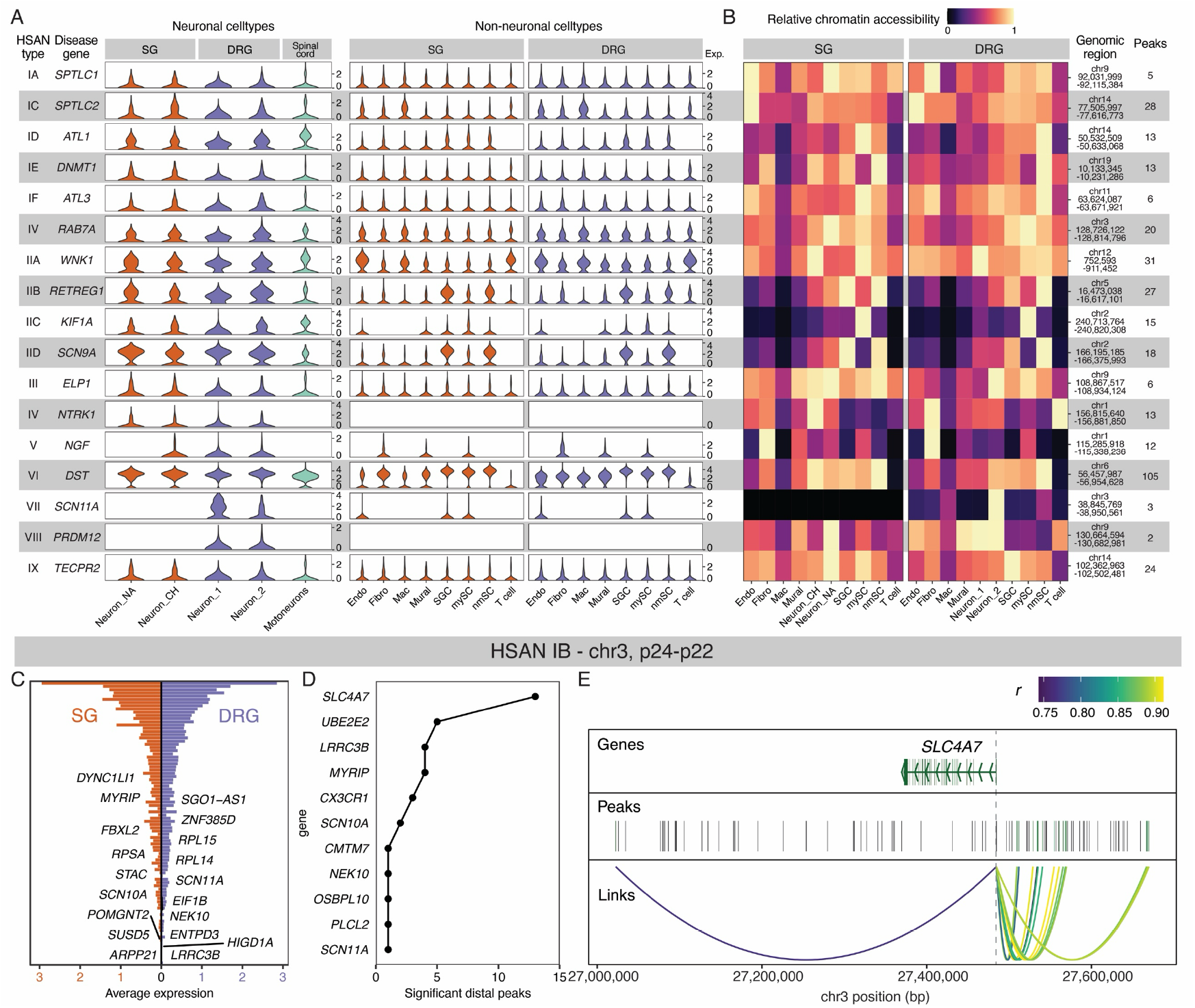
Cell types and genes implicated in hereditary sensory and autonomic neuropathies. **A**. Violin plots showing the expression of HSAN-associated genes in individual cell types from human SG and DRG, as well as the human spinal cord motoneurons previously reported. **B**. Heatmap showing the accessibility of genomic loci of HSAN-associated genes in individual human SG and DRG cell types. **C**. Bar plot showing the genes in the HSAN IB-associated locus that are also expressed in human SG and DRG. Genes that are highly expressed in neuronal cell types (Log2FC > 1, FDR < 0.05 comparing one neuronal cell type to all other nuclei) are highlighted. **D**. Genes that have significant associations with the distal peaks in the HSAN IB-associated locus. **E**. The correlation between snATAC-seq peaks with *SLC4A7* in the HSAN IB-associated locus. Distal peaks whose accessibility is highly correlated with *SLC4A7* expression in human SG and DRG cell types are highlighted and linked. SG: sympathetic ganglia, DRG: dorsal root ganglia. Endo: endothelial cell, Fibro: fibroblast, Mac: macrophage, SGC: satellite glial cell, mySC: myelinating Schwann cell, nmSC: non-myelinating Schwann cell. Neuron_NA: noradrenergic neuron, Neuron_CH: cholinergic neuron. Neuron_1: largely C-fiber neuron, Neuron_1: largely A-fiber neuron.

We first examined the expression pattern of HSAN-associated genes in human SG and DRG cell types from our atlas, as well as the motoneurons, whose axons innervate and terminate in the periphery, from a human spinal cord atlas previously reported[93]. Notably, genes associated with HSAN I subtypes (*SPTLC1, SPTLC2, ATL1, ATL3*, and *DNMT1*) and HSAN II subtypes (*WNK1, RETREG1, KIF1A*, and *SCN9A*) are expressed in all neuronal cell types, including those from SG and DRG, as well as motoneurons (Figure 6A), suggesting the mutant genes may directly and broadly affect the neuronal functions, corroborating the clinical observation of broad sensory, autonomic, and motor deficits in the patients. HSAN I and II-associated genes are also ubiquitously expressed in most non-neuronal cells from human SG and DRG, prompting the possibility that altered non-neuronal cell function may indirectly contribute to the axonal loss in HSAN I and II. The genomic loci of HSAN I and II-associated genes are broadly accessible in snATAC-seq data, consistent with the gene expression data (Figure 6B). Genes associated with HSAN III-VI (*ELP1, NTRK1, NGF*, and *DST*) and IX (*TECPR2*) are expressed, and their genomic loci are accessible in several neuronal and non-neuronal cell types from human SG and DRG (Figure 6A,B), consistent with the sensory and autonomic dysregulation reported in the patients. *SCN11A* is crucial for pain perception, and its mutations have been associated with HSAN VII[94, 95]. *SCN11A* is expressed 2.7 times more highly in C-fiber DRG neuron_1 cluster compared to A-fiber DRG neuron_2 cluster (Figure 6A), which may explain the patient’s inability to experience pain, whereas their sensitivity to temperature and touch are less affected. HSAN VII patients also experience mild autonomic dysfunctions, which could be mediated by the disruption of *SCN11A* normal function in the SG cells (Figure 6A), or the parasympathetic neurons that are not investigated in this study. *PRDM12* encodes a transcriptional regulator crucial for the development of nociceptors and HSAN VIII patients present with a profound insensitivity to pain [96, 97]. *PRDM12* is highly expressed in DRG neurons, and significantly higher in the C-fiber DRG neuron_1 cluster than the A-fiber DRG neuron_2 cluster, which is consistent with the clinical observation of insensitivity to pain. While *PRDM12* expression is only expressed in less than 1% of SG neurons (Figure 6A), the chromatin around its locus is accessible in SG noradrenergic neurons (Figure 6B). Future studies are warranted to understand the functions of *PRDM12* in the sympathetic neurons during development and how HSAN VIII-associated *PRDM12* mutations contribute to the autonomic deficits reported in the patients.

### Potential genes and GREs associated with HSAN IB

HSAN IB (also known as HSAN I with cough and gastroesophageal reflux) genome screen showed linkage to p24-p22 on chromosome 3 (3p24-22)[98, 99], but its exact mutations remain unknown. We reasoned that a better understanding of the functional genome of the human SG and DRG would help identify the possible genetic underpinnings of HSAN IB.

We first hypothesized that if the mutation(s) occur in the coding region of the genome, the expression of mutant gene(s) in the SG and DRG may contribute to the sensory and autonomic symptoms in HSAN IB patients. Therefore, we turned to our snRNA-seq data to identify the genes expressed in human peripheral ganglia that may be affected in HSAN IB. Out of 304 protein-coding genes annotated in 3p24-22, 88 are expressed (> 0.1 average counts) in at least one cell type (Figure 6C, S11), including 19 genes that are more highly expressed in neuronal cell types compared to non-neuronal cell types (Log2FC > 1, FDR < 0.05). *EIF1B* (Eukaryotic Translation Initiation Factor 1B), *RPL14, RPL15*, and *RPSA* (Ribosomal Proteins L14, L15, and SA, respectively) play critical roles in protein synthesis. *MYRIP* (Myosin VIIA and Rab Interacting Protein) and *DYNC1LI1* (Dynein Cytoplasmic 1 Light Intermediate Chain 1) are important for intracellular and axonal transport. *STAC* (SH3 and Cysteine Rich Domain) has been shown in animal models to regulate the voltage response of calcium channels and the release of neuropeptides, whereas ion channels, *SCN10A* and *SCN11A*, are crucial for the transmission of nociceptive signals. Mutations in those genes may lead to abnormal functions of the sensory and autonomic neurons, which could contribute to the underlying pathology of HSAN IB. In addition, given the important role of glial cells and immune cells in supporting and modulating neuronal functions, abnormal gene functions in these cell types may be a contributor to HSAN IB deficits (Figure S11).

As perturbation of GRNs may also lead to abnormal gene expression and function, we alternatively hypothesized that mutations in GREs could be another possible genetic cause of HSAN IB. To study the GRNs regulating gene expression in human SG and DRG cell types that may be associated with HSAN IB, we linked the snATAC-seq peaks in the distal, non-coding genomic region to individual expressed genes in 3p24-22. We found 10 genes with significant association (absolute value of *r* > 0.5) with 36 distal peaks (Figure 6D). For example, *SLC4A7* encodes a sodium bicarbonate cotransporter protein, which plays a crucial role in regulating intracellular pH. *SLC4A7* expression is positively correlated with the accessibility of 12 upstream peaks and one downstream peak (Figure 6E), suggesting that those peaks may contain cis-acting GREs that regulate the expression of *SLC4A7* when accessible. In summary, we identified 88 expressed genes and 36 accessible putative GREs tin human SG and DRG cell types. These genes and putative GREs are functionally relevant to human SG and DRG, and their disruptions may lead to dysfunctions of SG and DRG cell types, which may underlie the HSAN IB pathology. Our multi-omic atlas of human peripheral ganglia provides an invaluable opportunity for examining the diverse cell types in the human PNS that may be affected in HSAN and may provide novel insight into understanding the neurobiological mechanism of HSAN and identifying novel targets for therapeutic interventions.

## DISCUSSION

In this study, we generated a comprehensive single-cell multi-omic atlas of human peripheral ganglia, including SG and DRG, using snMultiome-seq. By analyzing tissues from multiple donors, we delineated the transcriptional and epigenomic landscapes of various cell types within these peripheral ganglia. This high-resolution atlas reveals shared and distinct molecular features of human SG and DRG cell types, providing new insights into the function and regulation of the human PNS. Additionally, we explored the potential implications of these cell types in peripheral neuropathies, specifically focusing on HSAN. This resource offers valuable insights into the physiological functions of these ganglia and their potential alterations in human diseases, which may inspire future investigation of the disease mechanisms and aid the development of novel therapeutic strategies.

Our integrated analysis showed a high level of similarity between human SG and DRG in terms of cellular composition and cell-type-specific gene expression. Both ganglia share major cell types, including neurons, glial cells, endothelial cells, fibroblasts, and immune cells. The composition of cell types is similar between human SG and DRG in our atlas, and the majority of the cell-type-specific marker genes are expressed at a similar level between the two ganglia. These cellular and molecular similarities likely reflect the shared developmental origin and the molecular infrastructures required for the basic biological functions of these peripheral ganglia.

Despite the overall similarities, distinct transcriptional signatures were observed between SG and DRG cell types, likely reflecting their unique physiological roles. The SG plays a critical role in maintaining homeostasis and mediating stress responses, whereas the DRG is important for relaying the sensory information to the CNS. Their distinct functional roles are exemplified by the molecular profiles of the neurons. We identified two human SG neuronal populations (Figure 1). The SG noradrenergic neurons express noradrenergic markers, such as *TH* and *DBH*. While also expressing noradrenergic markers, the SG cholinergic neurons uniquely express cholinergic markers, including *ACHE* and *SLC18A3*. Recent animal studies have suggested specificity in autonomic system response, which is likely mediated by SG neuronal populations. For example, transcriptionally distinct subsets of SG noradrenergic neurons specifically innervate erector muscles and control nipple- and pilo-erection [30], whereas cholinergic SG neurons innervate sweat glands and control sweat secretion[100, 101]. On the other hand, DRG neurons use glutamate to communicate with post-synaptic spinal neurons. A-fiber neurons are large-diameter, fast-conducting, and tend to be preferentially specialized for detecting mechanical stimuli, whereas C-fiber neurons are unmyelinated, slow-conducting, and can detect a broader range of environmental stimuli, including thermal, mechanical, and chemical stimuli[27, 102]. Compared to SG, DRG neurons highly express genes encoding ion channels for nociceptive signal transmission (e.g., *SCN10A* and *SCN11A*) and temperature sensing (*TRPA1* and *TRPM8*, Figure 3). These findings are consistent with previous literature.

Our cross-species comparison between human and mouse SG datasets also revealed some interesting differences. For example, we found proportionally more abundant myelinated, cholinergic neurons and mySCs in our human atlas, compared to the previous animal study[4]. While multiple possible factors may contribute to this difference (e.g., species difference and/or spinal-level difference), previous studies have suggested an expansion of cholinergic neurons in humans. In human SG, cholinergic neurons make up 18-28% of total human SG neurons in a previous study[40], and 33% of total human SG neurons in our atlas. In rodents, cholinergic neurons are estimated to comprise 5% of the neurons in the stellate SG, and as low as 0.1% in paravertebral SG[61, 62]. Physiologically, humans cool down body temperature by sweating, whereas other mammals have mostly hairy skins and generally dissipate heat in alternative ways, such as panting, increased saliva secretion, and blood flow in hairless skin[103]. Sweat glands, innervated and regulated by SG cholinergic neurons, are more prevalent in humans than in other mammals. These species differences in SG morphology and physiology previously described corroborate our observation of the expansion of human cholinergic neurons and mySCs. Therefore, our findings may reflect the physiological difference of SG innervation and the neurobiological basis of thermoregulation strategies across species.

Autonomic and sensory deficits have been reported in various human diseases, suggesting a potential perturbation of SG and DRG in these conditions. As the mechanisms by which the dysfunction of human SG and DRG may lead to the autonomic and sensory symptoms in human patients remain unclear, a better understanding of the molecular and cellular diversity of human SG and DRG may help us identify the cell types susceptible to these conditions and explore the potential molecular and cellular mechanisms of human diseases. For example, the accumulation of amyloid plaques, which usually happens when APP is erroneously processed, is a hallmark of Alzheimer’s disease. Amyloid plaques have been shown to reduce the neurite outgrowth of the sympathetic neurons in a mouse model of Alzheimer’s disease[56]. While autonomic dysfunctions are common in dementia patients[59, 60], their underlying neurobiological mechanisms have not been elucidated. Our analysis showed that *APP* is expressed in several SG cell types, which may contribute to the amyloid plaque accumulation and autonomic symptoms associated with Alzheimer’s (Figure 2). In addition, we examined the expression of *CNTN1* and *LAMA2*, genetic causes of specific types of congenital muscular dystrophies, in human SG [57, 58]. The expression of the mutant genes in SG cell types likely leads to altered normal SG functions and causes autonomic deficits in human patients, such as gastroesophageal reflux and, in more severe cases, respiratory failure[104]. Moreover, our multi-omic atlas has significant implications for the cell types that may be affected in HSAN. By mapping the transcriptional and epigenomic landscapes of HSAN-associated genes, we identified specific cell types likely to be affected in these disorders. Altogether, our human SG and DRG atlas provides insight into the key molecules and cell types associated with human health and diseases, and sheds light on the evolutionary conservation that would aid the future investigation of human diseases using animal models. This atlas can guide future research into the pathophysiology of HSAN and other neuropathies, facilitating the development of targeted therapeutic interventions.

Despite the comprehensive analyses described in this study, several limitations should be acknowledged. First, the low neuronal proportion in human PNS tissues undermines the absolute presentation of neuronal nuclei in our atlas, potentially leading to an underestimation of neuronal diversity within the human SG and DRG. Future studies with improved representation of neuronal populations (e.g., targeted approaches to enrich neurons and increased sample size) are necessary to capture the full extent of human neuron heterogeneity[27, 39, 42]. In addition, our atlas was generated using human tissues from organ donors, whose demographics may not be enriched for any specific diseases. While our analysis is promising for predicting and prioritizing cell types that may be affected in human diseases, the direct applicability of our findings to pathological conditions is limited and requires further investigation and validation. Future work incorporating tissues from patients or techniques introducing disease-associated mutations to non-pathological human tissues will be crucial for validating and extending our findings.

## MATERIALS AND METHODS

### Human tissue procurement

Post-mortem human tissues were obtained from organ donors with full legal consent for use of tissue in research and in compliance with procedures approved by Mid-America Transplant. The Human Research Protection Office at Washington University in St. Louis provided an institutional review board waiver. Extraction and collection of post-mortem human SG and DRG tissues were performed as previously described in collaboration with Mid-America Transplant[43], with the following modifications and specifications. Thoracolumbar paravertebral SGs and DRGs (T11 - L5) were surgically removed from postmortem organ donors, within 1 - 3 hours of aortic cross-clamping. Extracted tissues were immediately placed in ice-cold, oxygenated N-methyl-D-glucamine (NMDG)-based artificial cerebrospinal fluid (aCSF; 93 mM NMDG, 2.5 mM KCl, 1.25 mM NaH_2_PO_4_, 30 mM NaHCO_3_, 20 mM HEPES, 25 mM glucose, 5 mM ascorbic acid, 2 mM thiourea, 3 mM Na+ pyruvate, 10 mM MgSO_4_, 0.5 mM CaCl_2_, 12 mM N-acetylcysteine; adjusted to pH 7.3 using NMDG or HCl, and 300 – 310 mOsm using H_2_O or sucrose) and transported to the lab. Tissues were inspected, and adjacent tissues were removed. Cleaned tissues were snap-frozen in a liquid nitrogen vapor-based CryoPod Carrier before being transferred to −80 °C for long-term storage. To prepare for sequencing or RNA-FISH, human SG and DRG tissues were embedded in Optimal Cutting Temperature (OCT) compound and sectioned on a cryostat. Several sections were mounted for morphology inspection first, and five to ten 100-μm sections were collected into a nuclease-free 1.5 ml tube for sequencing. 15-μm sections were mounted on glass slides for RNA-FISH. Sections were stored at −80 °C if not used immediately.

### Single-nuclei isolation and gradient centrifugation

Nuclear extraction was performed according to a protocol described previously[28]. Tissues were rinsed with PBS to remove OCT and minced with scissors. Minced tissues were resuspended in 1 ml of homogenization buffer (0.25 M sucrose, 25 mM KCl, 5 mM MgCl_2_, 10 mM Tris-HCl, pH 8.0, 5 μg/ml actinomycin, 1% BSA, and 0.08 U/ul RNase inhibitor, 0.01% NP40) and transferred to a 2 ml dounce homogenizer on ice. Samples were homogenized for 15 strokes with the loose pestle, followed by 15 additional strokes with the tight pestle. The tissue homogenate was then passed through a 50 µm filter and diluted 1:1 with working solution (50% iodixanol, 25 mM KCl, 5 mM MgCl_2_, and 10 mM Tris-HCl, pH 8.0). Nuclei were layered onto an iodixanol gradient after homogenization and ultracentrifuged at 8,000 × g for 18 minutes. The gradient centrifugation was used to remove cellular and axonal debris and retain neuronal nuclei from the sample. After ultracentrifugation, nuclei were collected between the 30 and 40% iodixanol layers and diluted with resuspension buffer (1 × PBS with 1% BSA and 0.08 U/ul RNase inhibitor). Nuclei were centrifuged at 500 × g for 10 min at 4 °C and resuspended in resuspension buffer with 5 ng/μl of 7-AAD. Nuclei were inspected on a hemocytometer, and concentration was calculated. A step-by-step nuclear extraction protocol is available on protocols.io: https://dx.doi.org/10.17504/protocols.io.5jyl8q4y8l2w/v1.

During the preparation of the library hSG-1(Table 1), fluorescence-activated cell/nucleus sorting (FACS) was carried out in order to remove cellular debris. Nuclei were counterstained with 7-AAD and sorted using a 100 mm nozzle and a flow rate of 3 on a BD FACSARIA II into a 1.5 mL microcentrifuge tube containing 15 ml of 1X PBS, 0.04% BSA, and 0.1 U/ul RNase inhibitor. However, only 0.01% of the nuclei from this library were assigned to the neuronal clusters, compared to 2.4 ± 0.1% from libraries that were prepared without FACS. We concluded that FACS negatively affected the neuronal coverage and thus prepared the remaining libraries without FACS.

### snMultiome-seq

Nuclei were diluted to target 10k nuclei recovery for each library. Diluted nuclei were further processed and prepared for sequencing according to the manufacturer’s manuals of 10X Genomics Chromium Single Cell Multiome Assay. Libraries were sequenced on an Illumina NovaSeq X Plus instrument with 150 cycles each for Read1 and Read2, targeting 50,000 paired reads/nucleus for snRNA-seq libraries and 25,000 paired reads/nucleus for snATAC-seq libraries. Raw sequencing data from individual libraries were processed using 10X Genomics cellranger-arc (V2.0.1) and mapped to human reference genome GRCh38. Data from all eight snMultiome-seq libraries were then merged using cellranger-arc aggr function.

### Data processing, quality control, and clustering

The aggregated gene-cell and peak-cell count matrices were loaded and processed using R (V4.4.1) packages Seurat (V5.1.0) and Signac (V1.13.0)[52, 105]. To be included in the analysis, nuclei were required to contain more than 500 unique genes, less than 15,000 UMIs, and fewer than 5% of the counts deriving from mitochondrial genes for the snRNA-seq data, as well as more than 500 but fewer than 100,000 fragments, nucleosomal signal score less than 2, and TSS enrichment score greater than 1 for the snATAC-seq data. In total, there were 61,981 nuclei that met these initial quality control criteria.

Raw gene counts from snRNA-seq were scaled to 10,000 transcripts per nucleus and log-transformed using NormalizeData() function to control the sequencing depth between nuclei. Counts were centered and scaled for each gene using ScaleData() function. Highly variable genes were identified using FindVariableFeatures(), and the top 20 principal components were retrieved with RunPCA() using default parameters. For dimension reduction and visualization, Uniform Manifold Approximation and Projection (UMAP) coordinates for snRNA-seq (RNA-UMAP) were calculated using RunUMAP(). Nuclei clustering was performed using FindClusters() based on the variable features from the top 20 principal components, with the resolution set at 0.6. The marker genes for each cluster were identified using FindAllMarkers(), comparing nuclei in one cluster to all other nuclei. Doublet or low-quality nuclei were identified if they met any of the following criteria: 1). Assigned to a cluster with no significantly enriched marker genes (log2FC > 1, FDR < 0.05); 2). Identified as multiplets using R package DoubletFinder (V2.0.6) with doublet expectation rate at 5%[106]; 3). Assigned to a cluster in which three or more mitochondrial genes were identified among the top 20 marker genes (sorted by log2FC); and 4). Assigned to a cluster in which marker genes for multiple cell types were significantly enriched (log2FC > 1, FDR < 0.05). See the next section for the marker genes used. After iterative rounds of clustering and removal of low-quality clusters, a total of 52,950 high-quality human SG nuclei were included in the final dataset.

Raw peak counts of the high-quality nuclei from snATAC-seq were subsequently processed. Term frequency-inverse document frequency normalization was performed using RunTFIDF() function, and variable peaks were identified using FindTopFeatures(). Dimension reduction was performed with singular value decomposition using RunSVD() function, and UMAP coordinates for snATAC-seq (ATAC-UMAP) were calculated using RunUMAP(). Nuclei clustering was performed using FindClusters() based on the top 20 dimensions with the resolution of 0.6. To control donor variabilities, donor-specific peaks were called on snATAC-seq libraries from each donor using CallPeaks(), a MACS2 wrapper function in Signac. The snATAC-seq library derived from a pool of tissues of three donors was treated as a single donor. Only snATAC-seq peaks, originally identified by cellranger-arc and included in the peak-cell count matrix, that overlap with donor-specific peaks in no fewer than four donors were considered as shared peaks across donors and included in the analysis. ATAC-UMAP coordinates were re-generated by running RunUMAP() on shared peaks.

### Cell type annotation

Transcriptional cell types of human SG were assigned to each snRNA-seq cluster based on the canonical marker genes previously reported. Specifically, neuronal clusters are annotated based on the expression of *SNAP25* and *SYT1* (Figure S1E). Non-neuronal clusters are annotated based on the lack of expression of *SNAP25* and *SYT1*. Cell types were annotated based on the expression of canonical marker genes shown in Figure 1C. To identify subtypes and/or states of macrophages, we performed sub-clustering of the macrophages from human SG and DRG, and annotated the macrophage states using canonical marker genes previously reported (Figure S10A).

### Integration with mouse SG snRNA-seq data

To harmonize the SG cell types across species, our human SG snRNA-seq data were integrated with mouse SG snRNA-seq data previously described[4]. Mouse gene names were converted to their human orthologs using the R orthology mapping package Orthology.eg.db (V3.21). The two datasets were then jointly clustered using the clustering approach described above. To facilitate accurate comparative analysis across species, nuclei of different species were then integrated using IntegrateLayers() with CCA integration algorithm. FindClusters() and RunUMAP() were then run on the top 20 components of the integrated CCA space. To establish correspondence between human and mouse SG cell types, which were generated independently by the clustering approach described above, or published previously, the mouse cell type labels were transferred to each human nucleus by assigning the label of the most abundant mouse cell type in each integrated cluster to the human nuclei in the same integrated cluster. Mouse Schwann cells were renamed to mySCs in the analysis, as they express mySC marker *Ncamp*.

### Integration with human DRG snMultiome-seq data

To harmonize the SG cell types between human SG and human DRG, our human SG snMultiome-seq data were integrated with the human DRG snMultiome-seq data described in our companion study[38]. The two datasets were first jointly clustered using the clustering approach described above. To facilitate accurate comparative analysis, the two datasets were then integrated using IntegrateLayers() with CCA integration algorithm. FindClusters() and RunUMAP() were then run on the top 20 components of the integrated CCA space. To establish correspondence between human SG and DRG cell types, which were generated independently by the clustering approach described above, or published previously, the human DRG cell type labels were transferred to each human SG nucleus by assigning the label of the most abundant DRG cell type in each integrated cluster to the SG nuclei in the same integrated cluster.

### Differential analyses of gene expression and chromatin accessibility

To identify marker genes and peaks that are enriched in each cell type or macrophage subtype, differential expression/accessibility analysis was performed using FindAllMarkers() in Seurat/Signac, comparing nuclei from one cell type to all other nuclei. Genes and peaks with Log2FC > 1 and FDR < 0.05 were reported, unless specified otherwise.

To identify genes and peaks that are differentially expressed/accessible between any two populations (e.g., species-specific genes between human and mouse SG, or ganglion-specific genes between human SG and DRG), pair-wise differential expression/accessibility analysis was performed using FindMarkers() in Seurat/Signac, comparing nuclei of a cell type from one population to nuclei of the same cell type from another population. Genes and peaks with Log2FC > 1 and FDR < 0.05 were reported for the first population, and genes and peaks with Log2FC < (−1) and FDR < 0.05 were reported for the second population.

To identify sex-specific gene expression in each human SG cell type, pseudobulk differential expression analysis was performed to control for technical variations or donor variations across biological samples of the same sex. Specifically, pseudobulk counts for each cell type and each donor were generated by aggregating counts from nuclei of the same cell type and donor using AggregateExpression() in Seurat. Differential expression analysis was done using DESeq2 (V1.44.0) in R by comparing counts between male and female donors. Genes with Log2FC > 1 and FDR < 0.05 were reported. The snMultiome-seq library derived from a pool of tissues of three donors was excluded from the sex-specific differential analysis. To identify peaks that are differentially accessible between sexes, pair-wise differential accessibility analysis was performed using FindMarkers() in Signac, comparing nuclei of one cell type from male donors to nuclei of the same cell type from female donors. Peaks with Log2FC > 1 and FDR < 0.05 were reported.

### Ligand receptor pair analysis

Ligand receptor pair analysis was done using CellChat (V2.1.2)[107]. Human SG and mouse SG snRNA-seq data were analyzed separately. The ligand-receptor interaction databases CellChatDB.mouse and CellChatDB.human were used for mouse and human datasets, respectively. To infer the cell-type-specific communications, we first identified differentially expressed signaling genes in individual cell types using identifyOverExpressedGenes(). Significant interactions between ligands and receptors were identified using identifyOverExpressedInteractions(). Cell-cell communications were calculated using computeCommunProb() and communications present in less than 10 cells were excluded. Finally, cell-cell communications at the significant pathway level were quantified using computeCommunProbPathway().

### Gene ontology (GO) analysis

GO analysis was performed using topGO (V2.40.0) in R. Marker genes for each cell type were used as the input gene list. For comparison, the background gene list included all genes that are expressed in at least 5% of the nuclei in the cell type being analyzed. Genes were annotated for their biological process and associated GO terms. Enrichment is defined as the number of annotated genes observed in the input list divided by the number of annotated genes expected from the background list. GO terms with at least 20 annotated genes and enrichment *p*-value < 0.05 were returned.

### Functional Inference of Gene Regulation (FigR) analysis

To infer the transcription regulatory networks in human peripheral ganglia, we ran FigR on human SG and DRG snMultiome-seq datasets separately[68]. We first ran runGenePeakcorr() to calculate the Pearson’s *r* between the expression of genes in snRNA-seq data and the accessibility of peaks in snATAC-seq data falling within a 50,000bp window around each gene. Genes that are significantly associated (*p* < 0.05) with at least seven peaks were kept. Regulation score for each gene-peak association was calculated using getDORCScores(). To identify GRNs and link TFs to their putative regulated genes, FigR was run with default parameters by calling runFigRGRN(). FigR links TFs to putative regulated genes based on the TF motif enrichment in snATAC-seq peaks and the association of snATAC-seq peaks to genes previously described. TF motif enrichment was computed using chromVAR [108]. The snATAC-seq peak by TF motif overlap annotation matrix was first generated using a list of human TF motif PWMs from the chromVARmotifs package, and used along with the snATAC-seq reads in peak-cell count matrix to generate accessibility Z-scores across all snATAC-seq nuclei. To identify cell-type-specific regulatory mechanisms, we reported GRNs that met the following criteria: 1). The TF gene is differentially expressed in any human SG or DRG cell types (Log2FC>1, FDR<0.05); and 2). The TF motif is significantly enriched in the human SG or DRG dataset, with its average FigR enrichment > 0 across all gene-peak associations. GRNs with their average FigR regulation score > 0.14 across all gene-peak associations are identified as transcription activators, and TFs with their average FigR regulation score < (−0.07) across all gene-peak associations are identified as transcription repressors.

### Analysis of HSAN-associated genes and GREs

Genes and genomic loci associated with HSAN subtypes were retrieved from an online depository (https://neuromuscular.wustl.edu/time/hsn.htm) and previous studies[86, 109]. Expression of HSAN-associated genes in motoneurons was retrieved from snRNA-seq data of the human spinal cord previously described[93]. Ensemble genomic coordinates of HSAN-associated genes were retrieved using biomaRt (V2.60.1). To identify genes that may be affected by HSAN IB, all protein-coding genes in the genomic locus 3p24-22 (chromosome 3, 16,300,000-43,600,000bp) were retrieved using biomaRt. Genes with average expression > 0.1 in any human SG or DRG cell types are identified as expressed. To study the GREs that may be affected by HSAN IB, we first identified distal snATAC-seq peaks that are present in the non-coding genomic region in 3p24-22. To link distal peaks to putative regulated genes, we calculated Pearson’s *r* between the expression of genes expressed in human SG or DRG and the chromatin accessibility of distal peaks falling within a 50,000bp window around each gene. Significantly associated peaks to each gene were called with *r* > 0.5 or < (−0.5).

### RNA-FISH

RNA-FISH experiments were performed according to the manufacturer’s instructions, using the RNAscope Fluorescent Multiplex kit (Advanced Cell Diagnostics (ACD)) for fresh frozen tissue, as previously described[44]. Briefly, human SG and DRG tissues were frozen in OCT and sectioned into 15 μm sections using a cryostat. RNAScope probes against the following genes were ordered from ACD and multiplexed: Hs-SNAP25-C1 (Cat No. 518851), Hs-PRX-C2 (Cat No. 423911-C2), and Hs-NRXN1-C3 (Cat No. 527151-C3). Probes of *SNAP25, NRXN1*, and *PRX* were conjugated with OPAL-520, 570, and 690 for detection and visualization, respectively. Following RNA-FISH, slides were mounted in mounting media containing DAPI. RNA-FISH slides were imaged using a 20X objective on a confocal microscope for analysis, or using a 63X oil immersion objective on selected ROIs for visualization.

### FISH quantification

Two SGs and two thoracolumbar paravertebral DRGs from two donors were used for RNA-FISH, one SG and one DRG per donor (Table 1). Two to three non-consecutive sections from each biological sample were stained and used for quantification. For each section, two ROIs around axon regions (axon-enriched ROIs, less than five neurons in a ROI of ∼800 μm [h] × ∼800 μm [w], with clear morphology of nerve bundles) and two ROIs around neuronal soma regions (neuron-enriched ROIs, more than 20 neurons in the ROI defined above) were imaged. Five neuron-enriched ROIs and five axon-enriched ROIs per biological sample, or a total of 40 ROIs, were analyzed using cellpose and SCAMPR (Figure S8B)[66, 67]. First, cellpose was used to segment non-neuronal cells based on DAPI signal. Cellpose uses a machine learning algorithm that correctly identifies non-neuronal cells, as more than 98% of manually segmented cells were also segmented by cellpose. However, neurons had very dimmed DAPI staining and were poorly segmented by cellpose. Cell masks with an area > 200 μm^2^ or < 30 μm^2^ were excluded from the analysis. The resulting cell masks were converted into FIJI-compatible ROIs for downstream processing. In SCAMPR, background subtraction was applied, and image-specific threshold values were calculated per channel to correct the difference in background signal intensity across different images. A gene-cell matrix representing the expression of individual genes was generated based on the percentage of pixels in each cell expressing the gene of interest (the pixel intensity is greater than the image-specific threshold value previously mentioned). *PRX*+ cells were identified as cells with *PRX* expression greater than the 50-percentile across all *PRX*-expressing cells. *NRXN1*+ cells were identified as cells with *NRXN1* expression greater than the 50-percentile across all *NRXN1*-expressing cells. Lipofuscin autofluorescence was identified as large globular structures that exhibited highly similar fluorescent patterns across all three channels.

### Statistical analysis and visualization

Statistical analyses, including the number of samples or cells (n) and *p*-values for each experiment, are noted in the figure legends. Statistics and visualization were performed using R (V4.4.1). Student’s t-tests, ANOVA tests, and post-hoc t-tests were performed using the R package stats (V4.2.2). Hypergeometric tests were used to test the significance of the overlap between two groups by calling phyper() function also in the R package stats (V4.2.2). Plots were generated using R packages Seurat (V5.1.0), Signac (V1.13.0), ggplot2 (V3.5.1), gplots (V3.1.3), and CellChat (V2.1.2). Figure 1A was generated using bioRender.

## Acknowledgements

First and foremost, we express our deepest gratitude to the organ donors and their families for their invaluable gifts that made this research possible. We thank the past and present members of the Gereau lab for their helpful comments. We thank the Mid-America Transplant for their collaboration in tissue procurement. We thank the Genome Technology Access Center (GTAC) at McDonnell Genome Institute at Washington University School of Medicine for help with snMultiome-seq library construction and sequencing. GTAC is partially supported by NCI Cancer Center Support Grant #P30 CA91842 to the Siteman Cancer Center from the National Center for Research Resources (NCRR), a component of the National Institutes of Health (NIH), and NIH Roadmap for Medical Research.

## Author contributions

L.Y., A.C., and R.W.G. designed the sequencing experiments in this study. L.Y. performed the sequencing experiments of human SG. L.Y., A.J.D., K.B., and P.M. performed the sequencing analysis. L.Y., A.J.D., and R.W.G. designed the RNA-FISH experiments in this study, and L.Y., A.D., and R.T. performed and analyzed the RNA-FISH experiments. A.J.D., J.D.R., J.Y., R.S., Z.B., M.P., J.M.M., P.G., and J.L. extracted and processed the human SG and DRG tissues. L.Y. and R.W.G. wrote the manuscript with feedback from all authors. B.A.C., G.Z., V.C., A.C., and R.W.G. acquired funding and supervised all aspects of the study.

## Data and materials availability

Raw and processed data of human SG snMultiome-seq experiments included in this study will be deposited to the NCBI Gene Expression (GEO) SRA with accession number GSEXXX. Processed data from this study are also available on SPARC with DOI: https://doi.org/10.26275/oqxx-si9h. snRNA-seq data from mouse SG previously reported are available on GEO: GSE175421[4]. snMultiome-seq data from human DRG reported in our companion study are available on SPARC with DOI: https://doi.org/10.26275/sojr-qhvm[38]. A companion website for browsing the human SG and DRG data is available: https://peripheral-ganglia-multiome.shinyapps.io/atlas/.

