## Supplementary figures for "A multi-omic atlas of human autonomic and sensory ganglia implicates cell types in peripheral neuropathies"

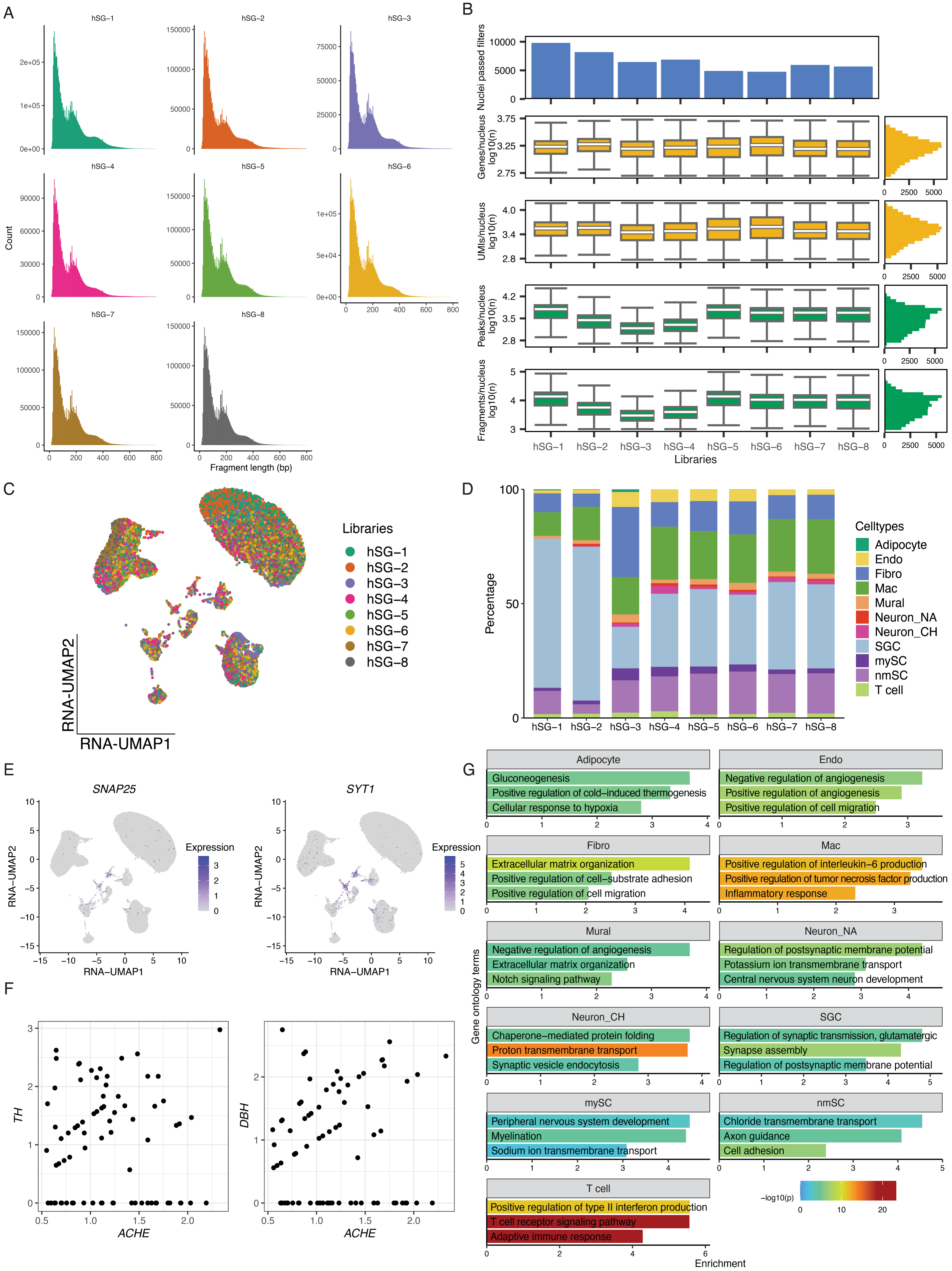
A
hSG−1
hSG−2
hSG−3
hSG−4
hSG−5
hSG−6
hSG−7
hSG−8
Count
Fragment length (bp)
B
Nuclei passed filters
Genes/nucleus log10(n)
UMIs/nucleus log10(n)
Peaks/nucleus log10(n)
Fragments/nucleus log10(n)
Libraries
C
Libraries
RNA-UMAP2
RNA-UMAP1
D
Percentage
Celltypes
Adipocyte
Endo
Fibro
Mac
Mural
Neuron_NA
Neuron_CH
SGC
mySC
nmSC
T cell
E
SNAP25
SYT1
Expression
RNA−UMAP2
RNA−UMAP1
F
TH
DBH
ACHE
G
Adipocyte
Gluconeogenesis
Positive regulation of cold−induced thermogenesis
Cellular response to hypoxia
Endo
Negative regulation of angiogenesis
Positive regulation of angiogenesis
Positive regulation of cell migration
Fibro
Extracellular matrix organization
Positive regulation of cell−substrate adhesion
Positive regulation of cell migration
Mac
Positive regulation of interleukin−6 production
Positive regulation of tumor necrosis factor production
Inflammatory response
Mural
Negative regulation of angiogenesis
Extracellular matrix organization
Notch signaling pathway
Neuron_NA
Regulation of postsynaptic membrane potential
Potassium ion transmembrane transport
Central nervous system neuron development
Neuron_CH
Chaperone−mediated protein folding
Proton transmembrane transport
Synaptic vesicle endocytosis
SGC
Regulation of synaptic transmission, glutamatergic
Synapse assembly
Regulation of postsynaptic membrane potential
mySC
Peripheral nervous system development
Myelination
Sodium ion transmembrane transport
nmSC
Chloride transmembrane transport
Axon guidance
Cell adhesion
T cell
Positive regulation of type II interferon production
T cell receptor signaling pathway
Adaptive immune response
Gene ontology terms
Enrichment
−log10(p)

**Figure S1 – Quality control metrics of human SG snMultiome-seq data, related to Figure 1**

**A**. Length distribution of snATAC-seq fragments mapped to the first 1M base pairs in Chromosome one in nuclei of each library. **B**. Quality control metrics of snMultiome-seq libraries. Top row displays the number of nuclei passed quality control. Second and third rows display the number of genes and UMIs per nucleus (log10 transformed) in snRNA-seq libraries. Bottom two rows display the number of peaks and transposase-sensitive DNA fragments per nucleus in snATAC-seq libraries. Boxes indicate quartiles and whiskers are 1.5-times the interquartile range. The median is a white line inside each box. The distribution is aggregated across all samples and displayed on the horizontal histogram. **C**. Gene expression UMAP of 10,000 randomly downsampled nuclei (1,250 nuclei per library). Nuclei were colored by libraries. **D**. Composition of transcriptional cell types in each snMultiome-seq library. **E**. Gene expression UMAP showing the expression of neuronal marker genes *SNAP25* (left) and *SYT1* (right). **F**. Scatter plots showing the co-expression of noradrenergic neuronal marker *ACHE* with cholinergic neuronal markers *TH* (left) or *DBH* (right) in individual nuclei from the cholinergic neuron cluster. **G**. Bar plot showing the top three significant gene ontology terms per cell type. Color denotes the p-values associated with each term.

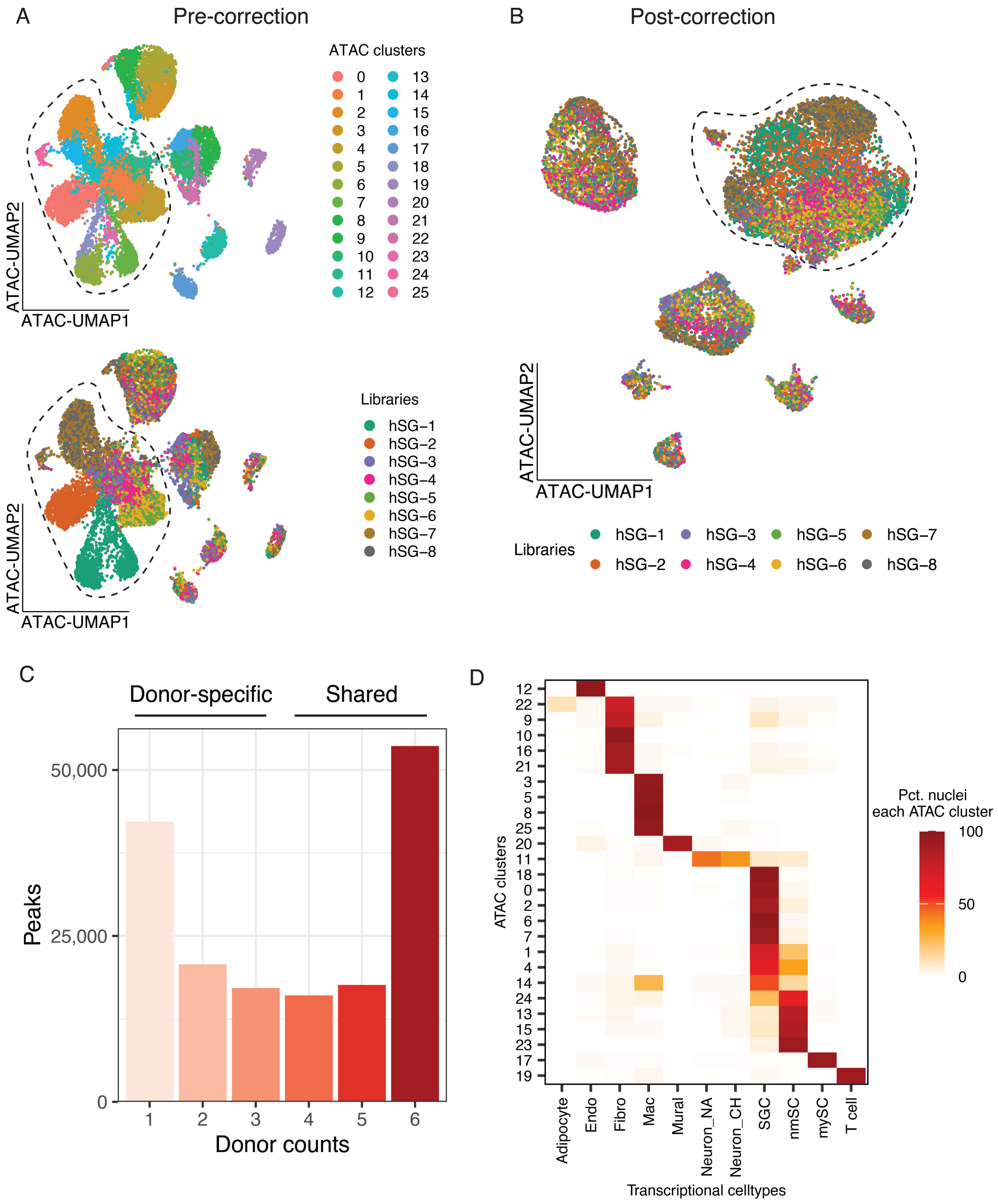


**Figure S2 – Integration of human SG snATAC-seq data, related to Figure 1**

**A**. Chromatin accessibility UMAP of 10,000 randomly downsampled nuclei (1,250 nuclei per library) clustered based on all peaks. Nuclei were colored by snATAC-seq clusters (top) or libraries (bottom). The dotted line highlights satellite glial cell clusters. **B**. Chromatin accessibility UMAP of 10,000 randomly downsampled nuclei (1,250 nuclei per library) clustered based on shared peaks defined in S2C. Nuclei were colored by libraries. The dotted line highlights satellite glial cell clusters. **C**. Distribution of snATAC-seq peaks based on their accessibility across donors. Peaks are called per donor using MACS2, and snATAC-seq peaks present in less than four donors are classified as 'donor-specific', and peaks present in at least four donors are classified as 'shared' and subsequently used in the snATAC-seq clustering in S2B. **D**. Overlap of human SG cell types between the snATAC-seq clustering and the transcriptional cell types assigned by snRNA-seq clustering. Plot displays the fraction of nuclei within each snATAC-seq cluster that is assigned to each transcriptional cell type.

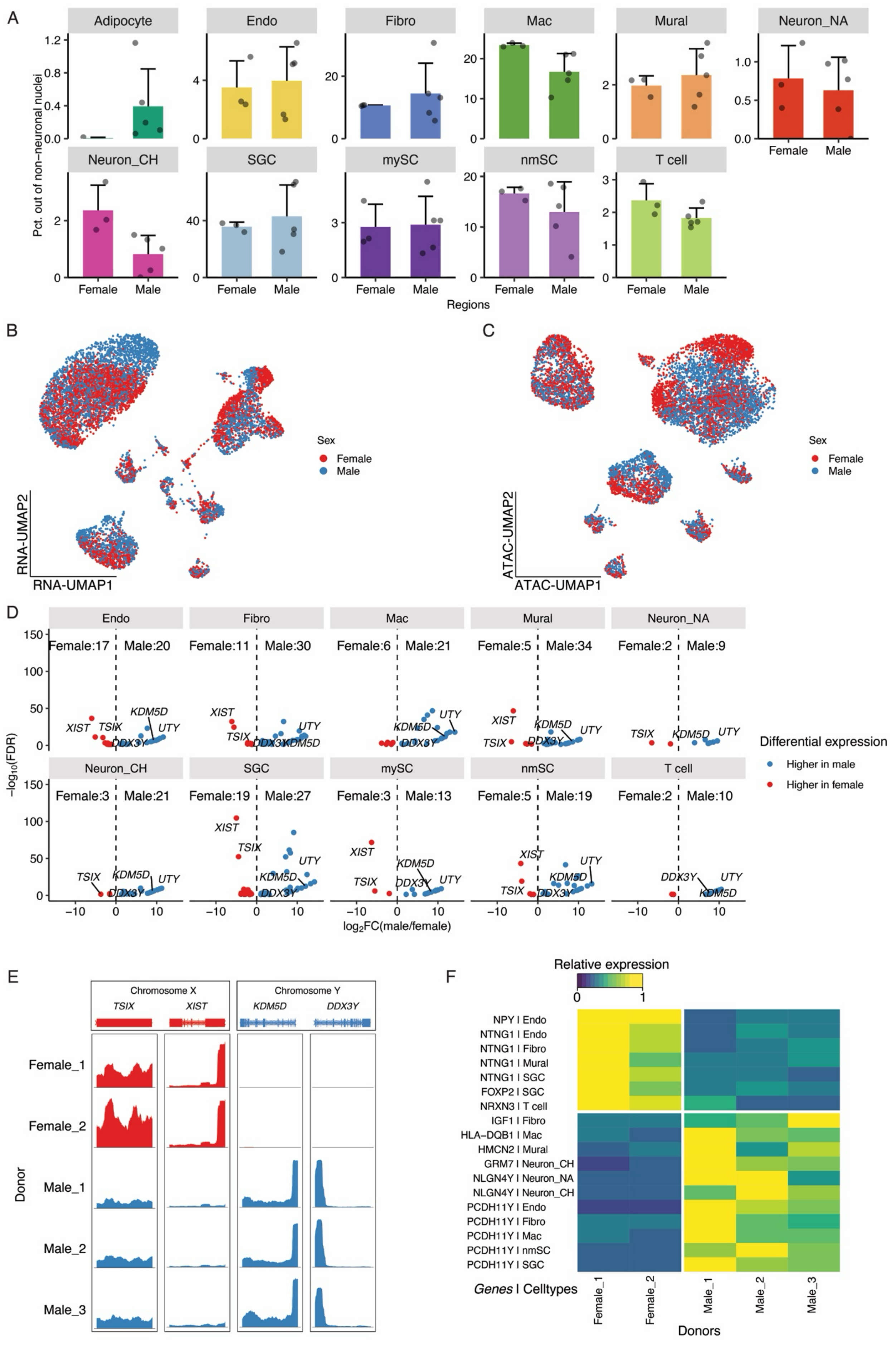

A
Adipocyte
Endo
Fibro
Mac
Mural
Neuron_NA
Neuron_CH
SGC
mySC
nmSC
T cell
Pct. out of non-neuronal nuclei
Female
Male
Regions
B
RNA-UMAP2
RNA-UMAP1
Sex
Female
Male
C
ATAC-UMAP2
ATAC-UMAP1
D
Female:17
Male:20
Female:11
Male:30
Female:6
Male:21
Female:5
Male:34
Female:2
Male:9
Female:3
Male:21
Female:19
Male:27
Female:3
Male:13
Female:5
Male:19
Female:2
Male:10
XIST
TSIX
KDM5D
UTY
DDX3Y
$-\log_{10}$(FDR)
$\log_2$FC(male/female)
Differential expression
Higher in male
Higher in female
E
Chromosome X
Chromosome Y
TSIX
XIST
KDM5D
DDX3Y
Female_1
Female_2
Male_1
Male_2
Male_3
Donor
F
Relative expression
NPY | Endo
NTNG1 | Endo
NTNG1 | Fibro
NTNG1 | Mural
NTNG1 | SGC
FOXP2 | SGC
NRXN3 | T cell
IGF1 | Fibro
HLA−DQB1 | Mac
HMCN2 | Mural
GRM7 | Neuron_CH
NLGN4Y | Neuron_NA
NLGN4Y | Neuron_CH
PCDH11Y | Endo
PCDH11Y | Fibro
PCDH11Y | Mac
PCDH11Y | nmSC
PCDH11Y | SGC
Genes | Celltypes
Female_1
Female_2
Male_1
Male_2
Male_3
Donors

**Figure S3 – Putative sex-specific transcriptional and epigenomic features, related to Figure 1**

**A**. Bar plot showing the composition of cell types in SG libraries between sexes. ($p = 0.99$ for adipocytes, $p = 0.89$ for endothelial cells, $p = 0.71$ for fibroblasts, $p = 0.03$ for macrophages, $p = 0.98$ for mural cells, $p = 0.99$ for noradrenergic neurons, $p = 0.71$ for cholinergic neurons, $p = 0.9$ for satellite glial cells, $p = 0.99$ for myelinating Schwann cells, $p = 0.99$ for non-myelinating Schwann cells, and $p = 1$ for T cells. Bonferroni-corrected student t-tests [*$p = 0.0045$, **$p = 0.0009$, ***$p = 0.00009$]. **B** and **C**. Gene expression (B) and chromatin accessibility (C) UMAPs of 10,000 randomly downsampled nuclei (5,000 nuclei per sex). Nuclei were colored by donors' sex. **D.** Volcano plots showing the differentially expressed genes (Log2FC > 1 for male and Log2FC < (-1) for female, FDR < 0.05) in individual cell types between nuclei from male donors and female donors. Known sex-specific genes were labeled. Cell types with less than 50 nuclei per sex were excluded from the analysis. **E**. Coverage plot showing the accessibility around genomic loci of known sex-specific genes in snATAC-seq libraries from individual donors. The chromatin accessibility is displayed as the relative frequency of sequenced DNA fragments for each donor, grouped by 50 bins per displayed genomic region. The frequency is normalized by the maximal frequency per genomic region. **F**. Heatmap showing the relative expression of selected sex-specific genes in a given cell type in male and female donors. Color denotes relative expression of a gene in the cell type specified across all donors displayed (calculated as the mean expression of a gene relative to the highest mean expression of that gene across all donors displayed).

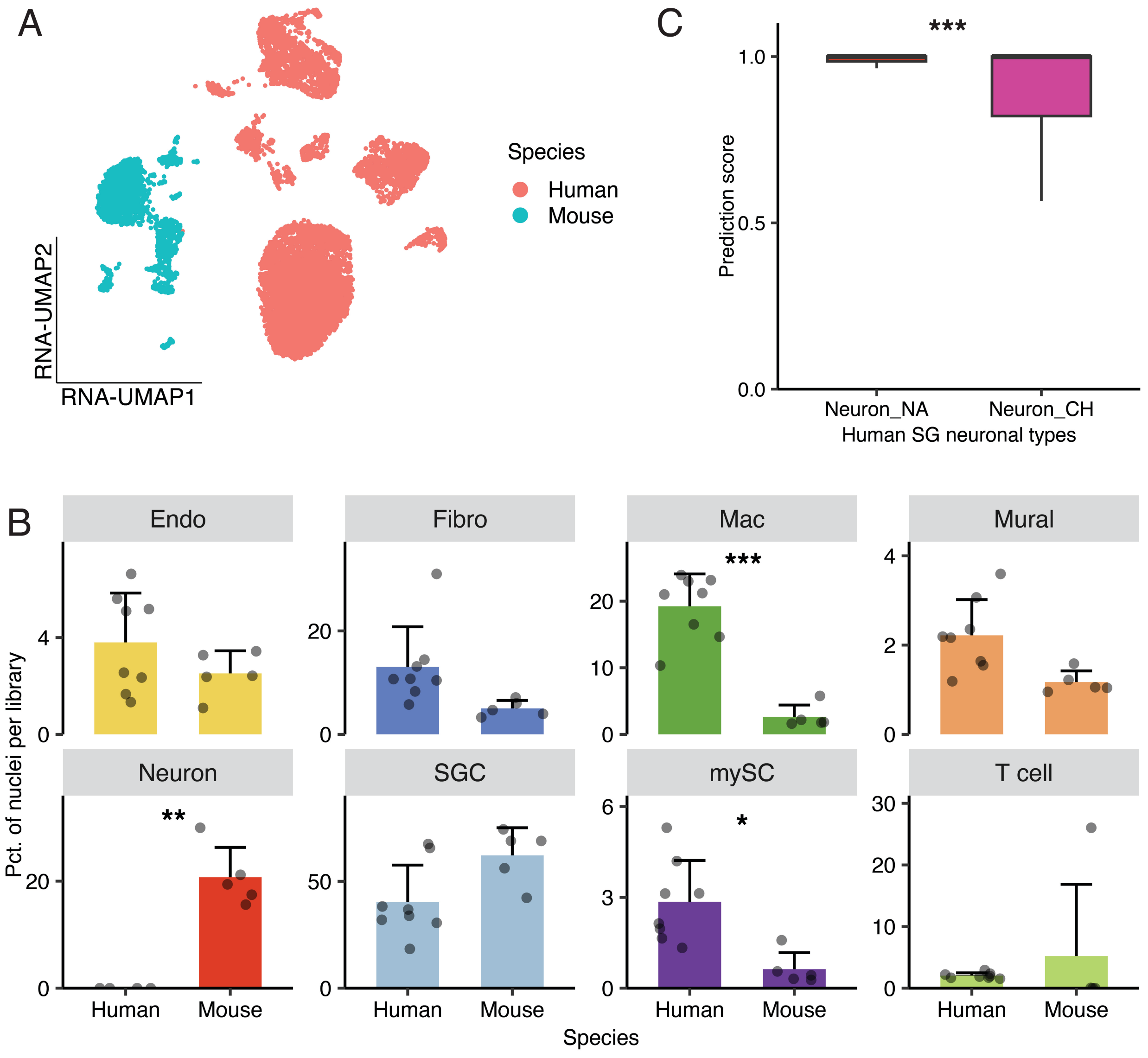


**Figure S4 – Comparison between human and mouse SG, related to Figure 2**

**A**. Gene expression UMAP of human and mouse SG when clustering together without integration. 10,000 nuclei (5,000 nuclei per species) were randomly sampled. Nuclei were colored by species. **B**. Bar plots showing the percentage of cell types in human and mouse SG libraries. ($p = 0.15$ for endothelial cells, $p = 0.02$ for fibroblasts, $p = 7.4e\text{-}6$ [***] for macrophages, $p = 7.4e\text{-}3$ for mural cells, $p = 1.1e\text{-}3$ [**] for neurons, $p = 0.02$ for satellite glial cells, $p = 2.1e\text{-}3$[*] for myelinating Schwann cells, and $p = 0.57$ for T cells. Bonferroni-corrected student t-tests comparing percentages of a given cell type in human libraries to those in mouse libraries [*$p = 6.3e\text{-}3$, **$p = 1.3e\text{-}3$, ***$p = 1.3e\text{-}4$]. **C**. Box plot showing the prediction score when anchoring human neuronal nuclei, including both noradrenergic and cholinergic clusters, to the mouse neuronal nuclei, which are mostly noradrenergic. ($p = 2.2e\text{-}16$ [***], student t-test).

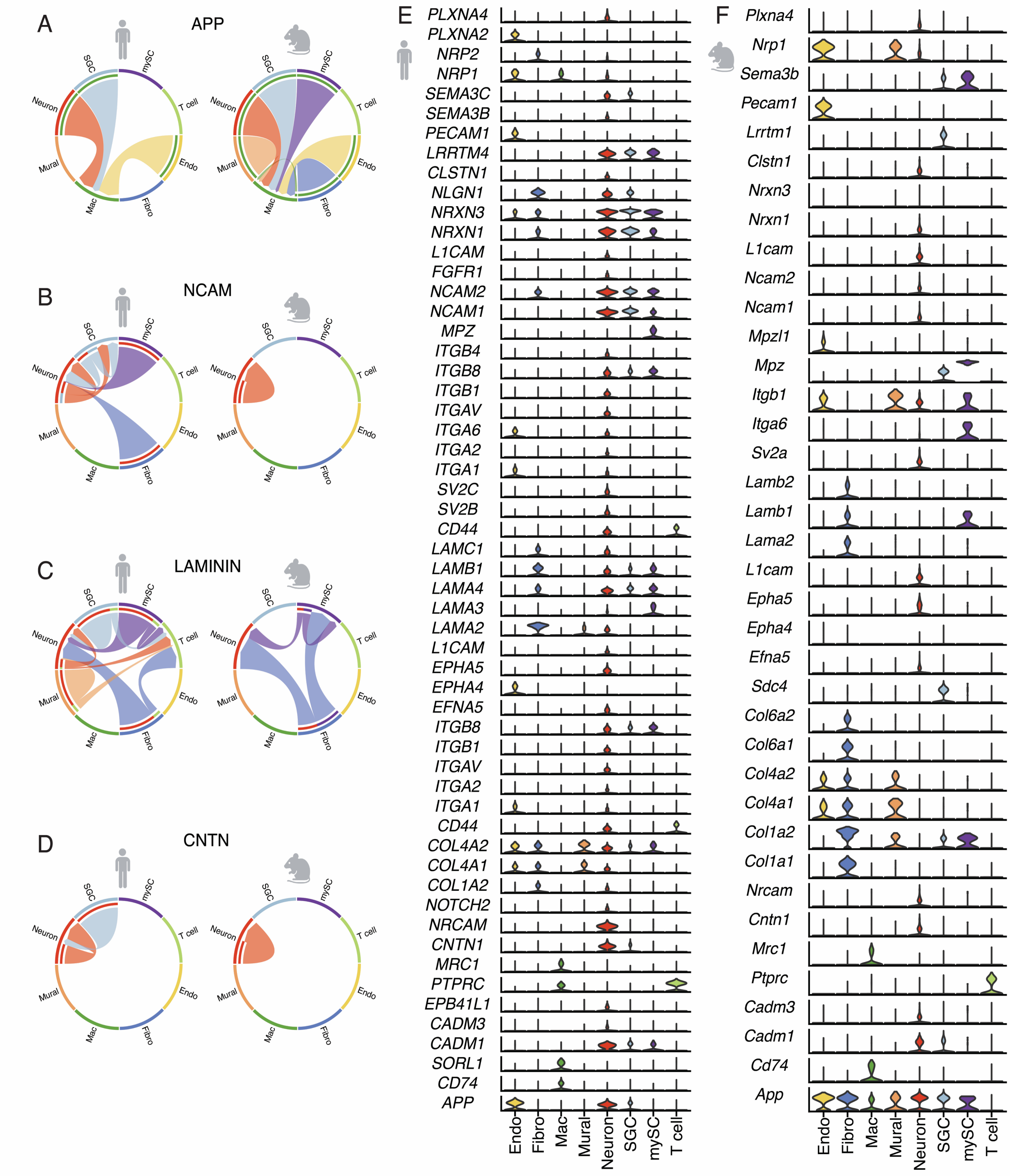


**Figure S5 – Conserved signaling pathways between human and mouse SG, related to Figure 2**

**A-D**. Chord diagrams showing selected signaling pathways significantly enriched in both human (left) and mouse (right) SG data. In the chord diagram, the inner thinner bar colors represent the targets that receive signals from the corresponding outer bar. The inner bar size is proportional to the signal strength received by the targets. **E** and **F**. Violin plots showing the expression level across cell types of genes associated with the conserved signaling pathways in human (E) and mouse (F). Violin plots were colored by cell types. Human nuclei from noradrenergic and cholinergic clusters were merged as one neuronal cell type for ligand receptor analysis.

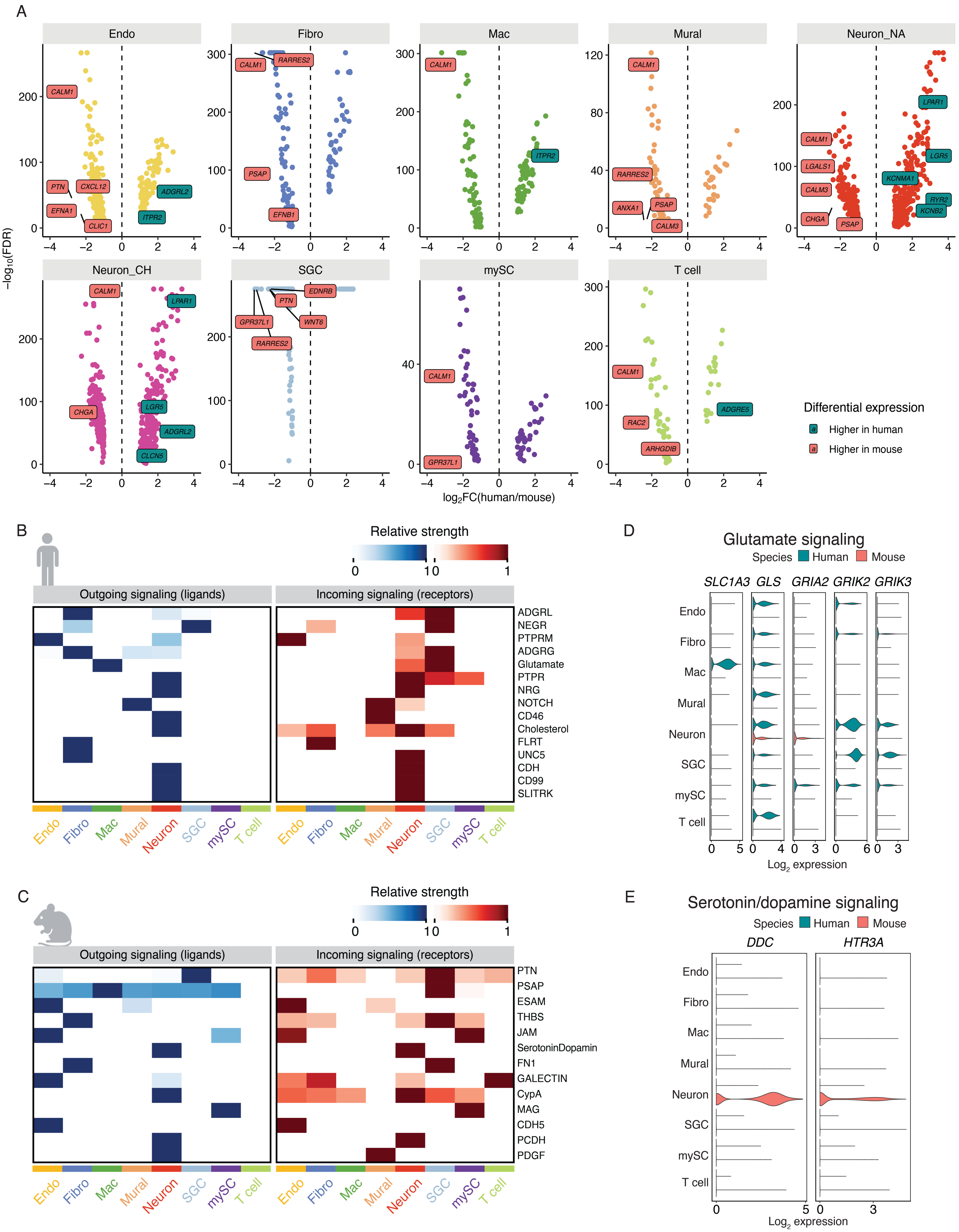
A
Endo
Fibro
Mac
Mural
Neuron_NA
Neuron_CH
SGC
mySC
T cell
−log10(FDR)
log2FC(human/mouse)
CALM1
PTN
CXCL12
EFNA1
CLIC1
ADGRL2
ITPR2
RARRES2
PSAP
EFNB1
ANXA1
CALM3
LGALS1
CHGA
LPAR1
LGR5
KCNMA1
RYR2
KCNB2
CLCN5
EDNRB
WNT6
GPR37L1
RAC2
ARHGDIB
ADGRE5
Differential expression
Higher in human
Higher in mouse
B
Relative strength
Outgoing signaling (ligands)
Incoming signaling (receptors)
ADGRL
NEGR
PTPRM
ADGRG
Glutamate
PTPR
NRG
NOTCH
CD46
Cholesterol
FLRT
UNC5
CDH
CD99
SLITRK
Neuron
C
PTN
PSAP
ESAM
THBS
JAM
SerotoninDopamin
FN1
GALECTIN
CypA
MAG
CDH5
PCDH
PDGF
D
Glutamate signaling
Species
Human
Mouse
SLC1A3
GLS
GRIA2
GRIK2
GRIK3
Log2 expression
E
Serotonin/dopamine signaling
DDC
HTR3A

**Figure S6 – species-specific features between human and mouse SG, related to Figure 2**

**A**. Volcano plots showing the differentially expressed genes (Log2FC > 1 for human and Log2FC < (-1) for mouse, FDR < 0.05) in individual cell types between human and mouse snRNA-seq data. Top 5 genes (sorted by Log2FC) of neuropeptides, neuropeptide receptors, GPCR, and ion channels per cell type and species were labeled. **B** and **C**. Heatmaps showing the relative strength of signaling pathways enriched only in human (B) or mouse (C) data. Outgoing signaling strength is calculated by aggregating the average expression of ligands associated with a given pathway in each cell type. Incoming signaling strength is calculated by aggregating the average expression of receptors associated with a given pathway in each cell type. The strength of a signal pathway is normalized by the maximal strength across all cell types in a given species. **D** and **E**. Violin plots showing the expression of genes associated with the glutamate signaling pathway (D) and serotonin/dopamine signaling pathway (E) in individual human and mouse cell types. Violin plots were colored by species. Human nuclei from noradrenergic and cholinergic clusters were merged as one neuronal cell type for ligand receptor analysis.

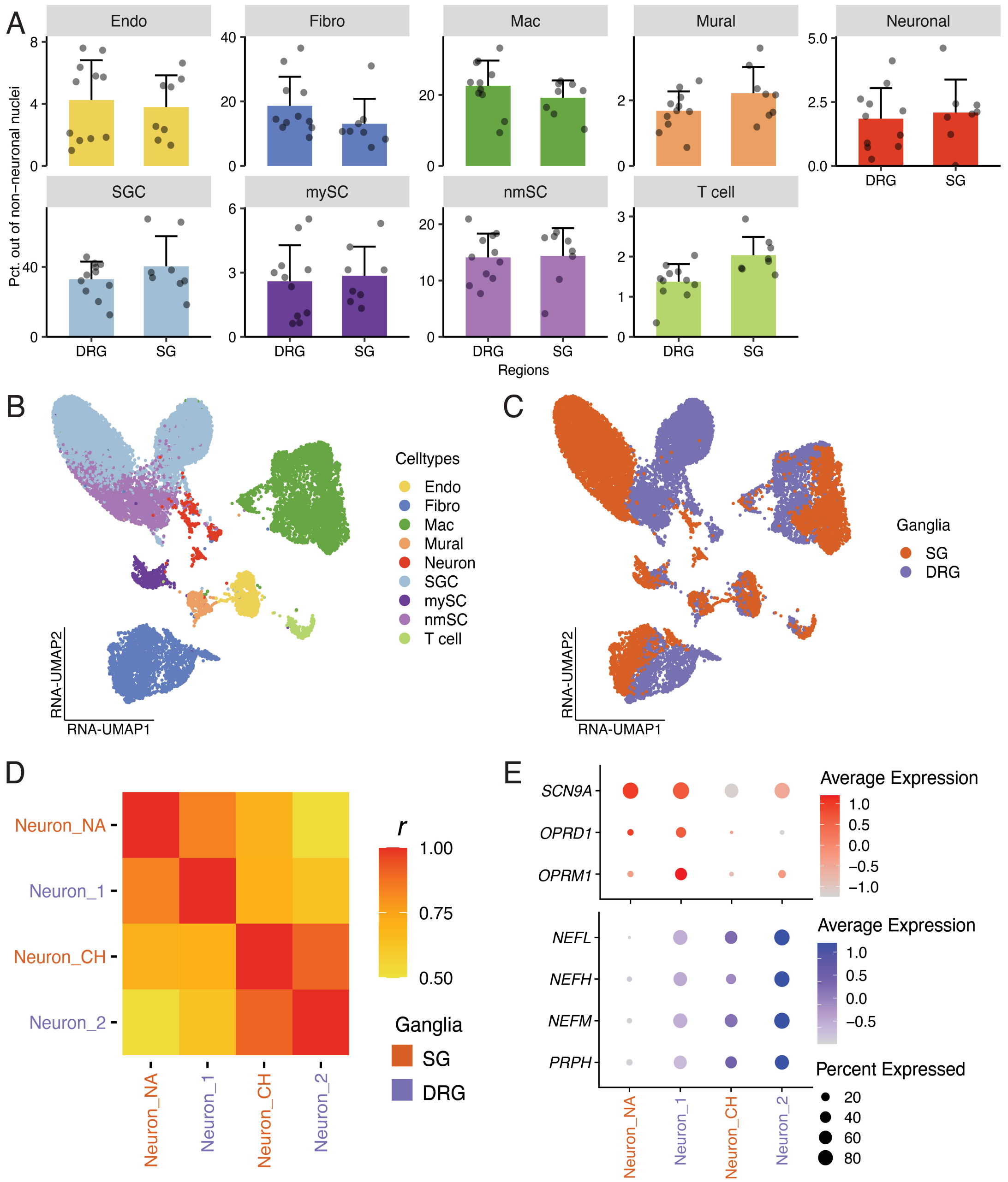


**Figure S7 – Comparison between human SG and DRG, related to Figure 3**

**A.** Bar plot showing the cell type composition in human SG and DRG libraries. ($p = 0.91$ for endothelial cells, $p = 0.08$ for fibroblasts, $p = 0.24$ for macrophages, $p = 0.96$ for mural cells, $p = 0.28$ for neuronal [all neuronal clusters combined], $p = 0.93$ for satellite glial cells, $p = 0.97$ for myelinating Schwann cells, $p = 0.3$ for non-myelinating Schwann cells, and $p = 1$ for T cells). Bonferroni-corrected student t-tests comparing percentages of a given cell type in human libraries to those in mouse libraries [*$p = 0.005$, **$p = 0.0011$, ***$p = 0.00011$]. **B and C.** Gene expression UMAP of human SG and DRG when clustering together without integration. 10,000 nuclei (5,000 nuclei per dataset) were randomly sampled. Nuclei were colored by transcriptional cell types (B) or ganglia (C). **D.** Heatmap showing the Pearson's correlation coefficient (*r*) by correlating the expression of cell-type-specific marker genes between human SG and DRG neuronal cell types. **E.** Dot plot showing the expression of selected genes shared between SG noradrenergic neurons and DRG neuron_2 cluster (genes colored red), or SG cholinergic neurons and DRG neuron_1 cluster (genes colored blue).

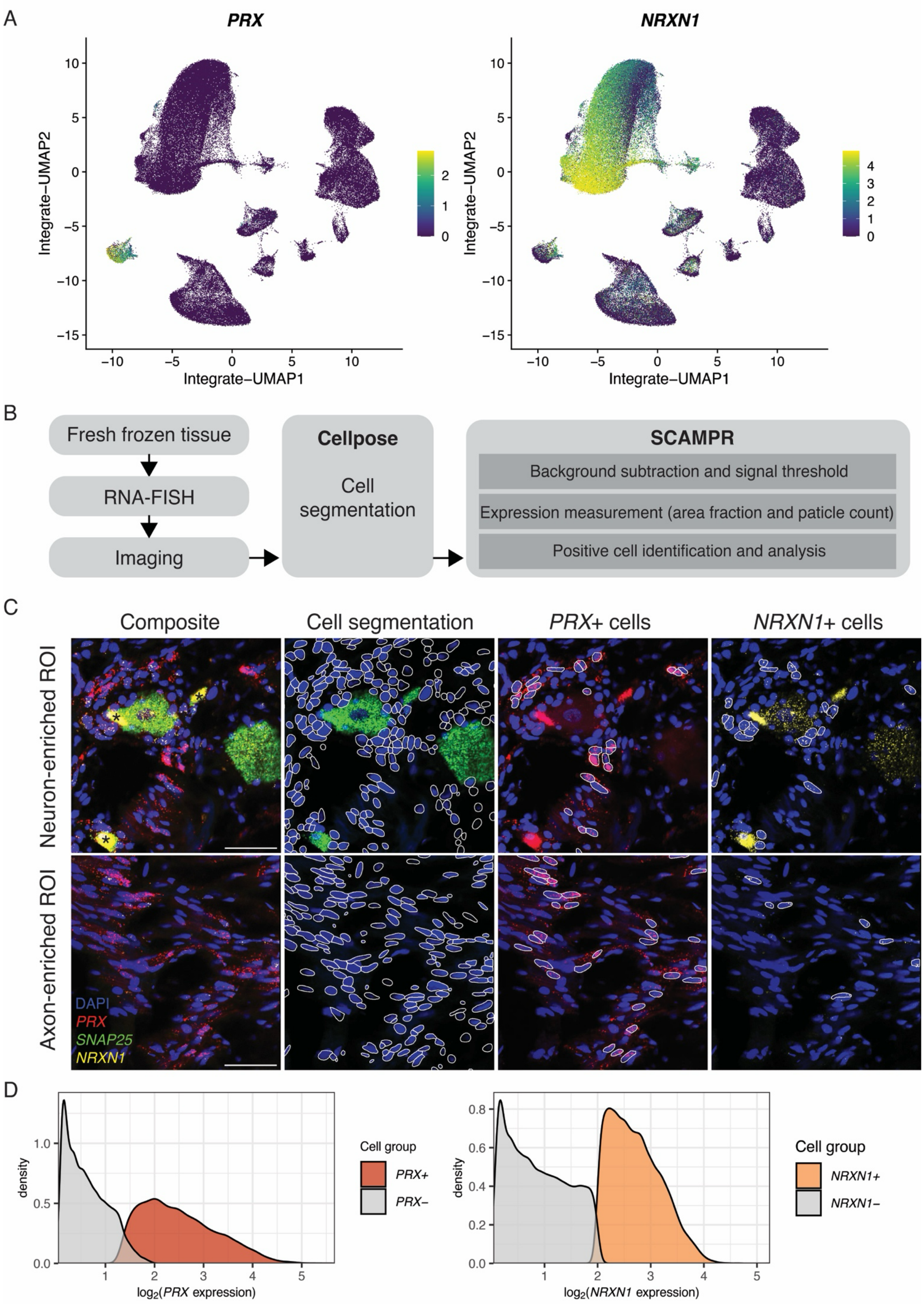

A
PRX
NRXN1
Integrate−UMAP1
Integrate−UMAP2
B
Fresh frozen tissue
RNA-FISH
Imaging
Cellpose
Cell segmentation
SCAMPR
Background subtraction and signal threshold
Expression measurement (area fraction and paticle count)
Positive cell identification and analysis
C
Composite
Cell segmentation
PRX+ cells
NRXN1+ cells
Neuron-enriched ROI
Axon-enriched ROI
DAPI
PRX
SNAP25
NRXN1
D
density
log2(PRX expression)
Cell group
PRX+
PRX−
log2(NRXN1 expression)
Cell group
NRXN1+
NRXN1−

**Figure S8 – Glial cell quantification in human SG and DRG, related to Figure 4**

**A.** Integrative gene expression UMAPs showing the expression of myelinating Schwann cell marker *PRX* (left) and satellite glial and non-myelinating Schwann cell marker *NRXN1* (right) in human SG and DRG snMultiome-seq data. **B.** Workflow of RNA-FISH image analysis. **C.** Representative images demonstrating results of cell segmentation and calling of positive cells based on *PRX* and *NRXN1* expression. Scale bar = 50 um. **D.** Distribution of *PRX* and *NRXN1* expression in all expressing cells.

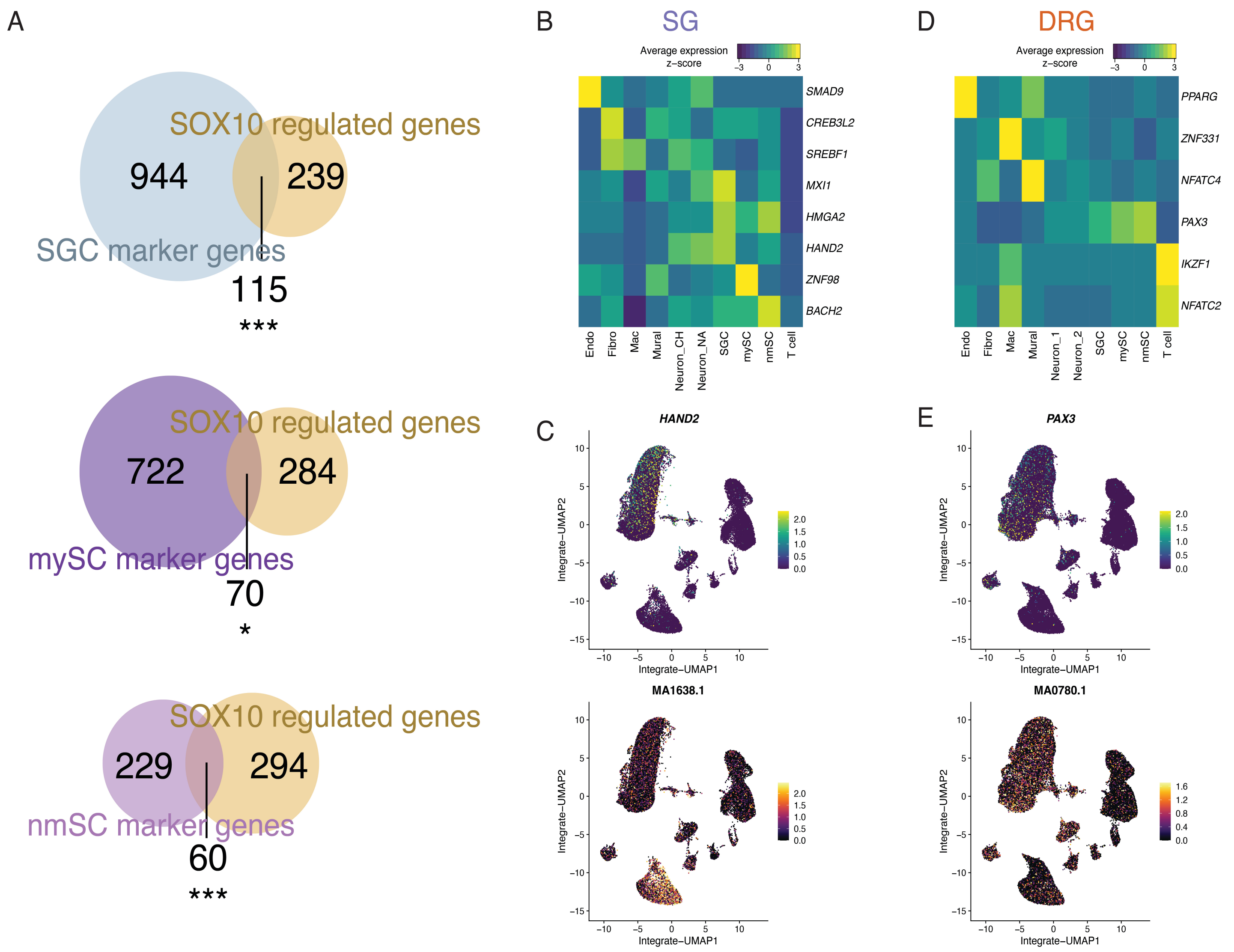


**Figure S9 – Genomic regulatory networks in human SG and DRG, related to Figure 5**

**A.** Overlap between SOX10-regulated genes with marker genes for satellite glial cells (top), myelinating Schwann cells (mid), and non-myelinating Schwann cells (bottom). Hypergeometric tests: $p$ = 2.4E-8 [***] for satellite glial cells, $p$ = 0.017 [*] for myelinating Schwann cells, $p$ = 1,2E-17 [***] for non-myelinating Schwann cells. [*$p$ = 0.05, ***$p$ = 0.001] **B.** Heatmap showing the expression of TFs significantly enriched only in SG data in individual SG cell types. **C.** UMAPs showing *HAND2* gene expression (top) in human SG snRNA-seq data and HAND2 motif enrichment (bottom) in human SG snATAC-seq data. **D.** Heatmap showing the expression of TFs significantly enriched only in DRG data in individual DRG cell types. **E.** UMAPs showing *PAX3* gene expression (top) in human DRG snRNA-seq data and PAX3 motif enrichment (bottom) in human DRG snATAC-seq data.

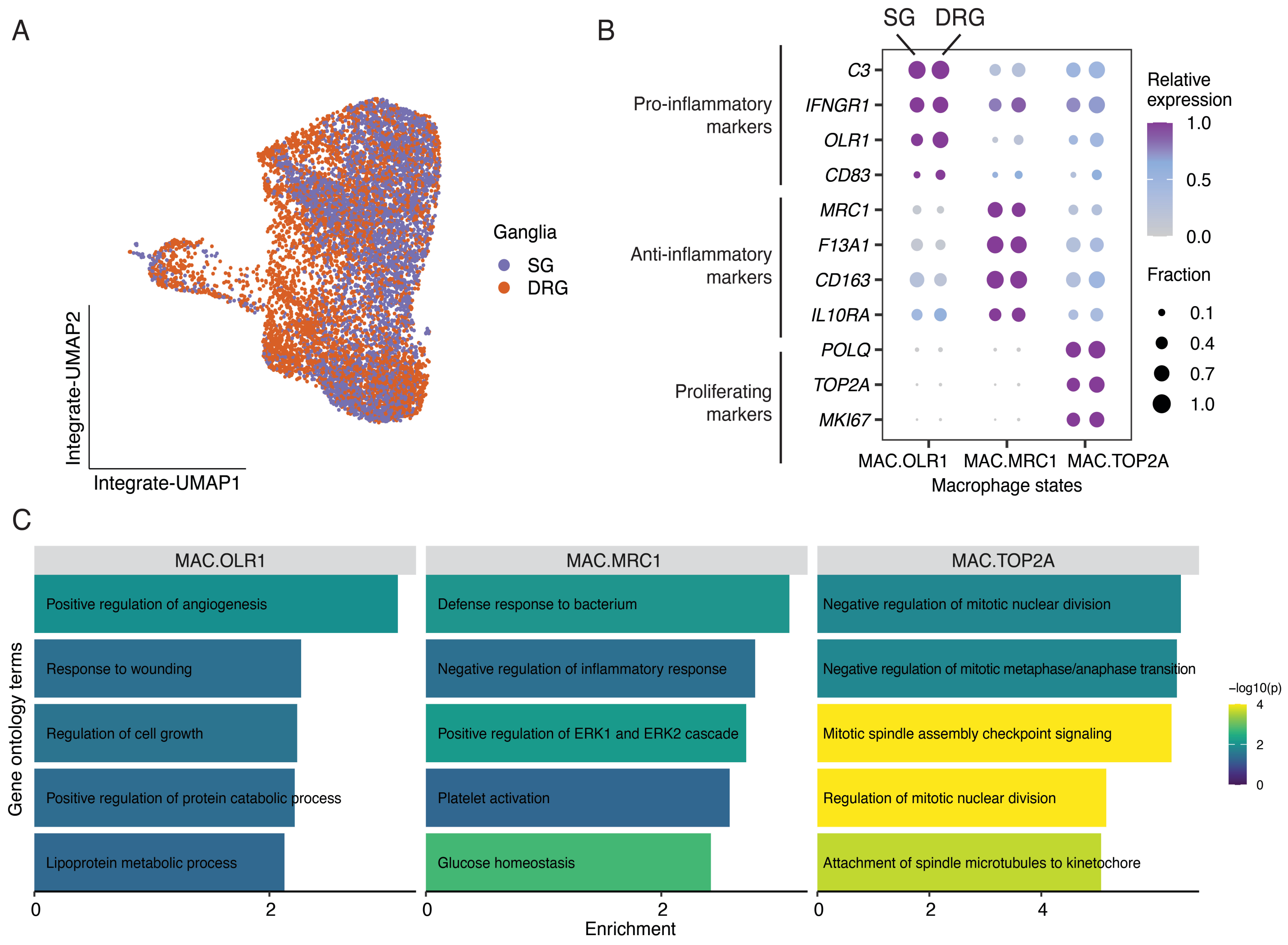


**Figure S10 – Subclustering of macrophages in human SG and DRG, related to Figure 5**

**A.** Integrative gene expression UMAP showing the subtypes of 10,000 randomly sampled (5,000 nuclei per ganglion) macrophage nuclei. Nuclei were colored by ganglia. **B.** Dot plot displaying the expression of marker genes in individual macrophage clusters/states. Dot size denotes the fraction of nuclei expressing a marker gene (>0 counts), and color denotes relative expression of a gene in each macrophage subtype (calculated as the mean expression of a gene relative to the highest mean expression of that gene across all cell types). **C.** Top five gene ontology terms enriched in each macrophage cluster.

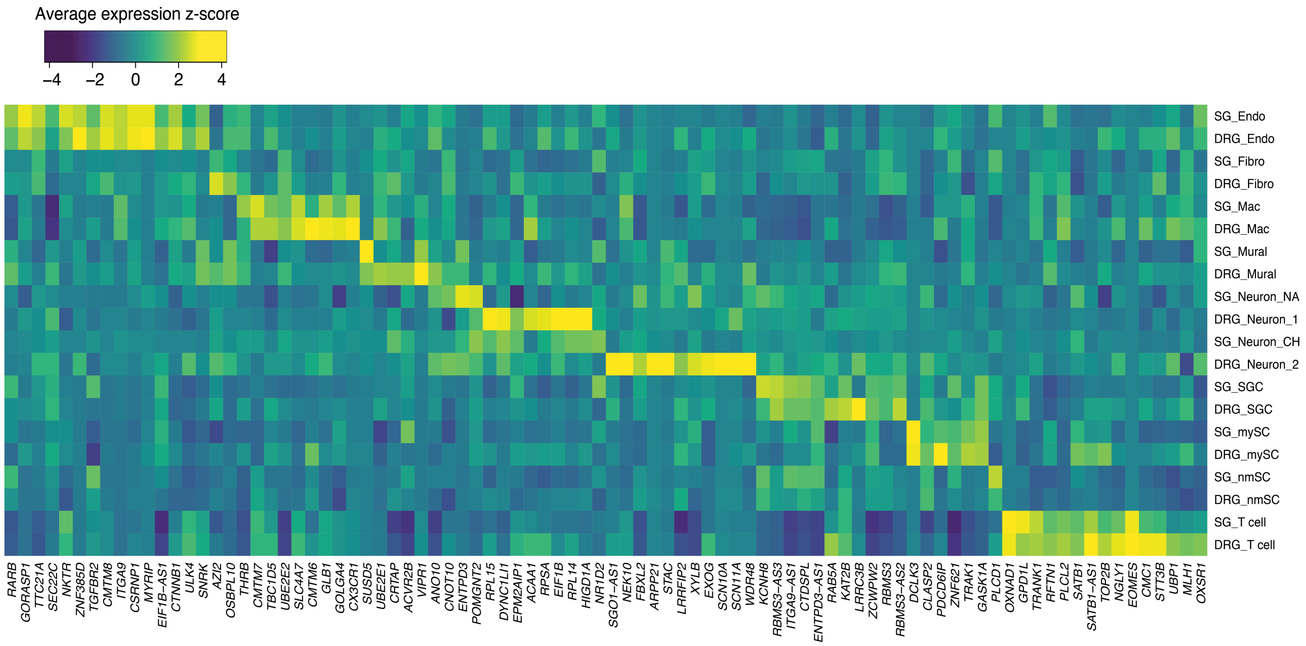


**Figure S11 – Genes in HSAN 1B-asscoaited locus, related to Figure 6**

Heatmap showing the expression of genes whose gene body overlaps with 3p24-22, that are also expressed in any cell types (expressed in at least 5% nuclei) in human SG and DRG snRNA-seq data.
